# FACT regulates pluripotency through distal regulation of gene expression in murine embryonic stem cells

**DOI:** 10.1101/2021.07.30.454509

**Authors:** David C. Klein, Santana M. Lardo, Kurtis N. McCannell, Sarah J. Hainer

## Abstract

The FACT complex is a conserved histone chaperone with essential roles in transcription and histone deposition. FACT is essential in pluripotent and cancer cells, but otherwise dispensable for most mammalian cell types. FACT deletion or inhibition can block induction of pluripotent stem cells, yet the mechanism through which FACT regulates cell fate decisions remains unclear. To determine this mechanism, we used inducible depletion of FACT subunit SPT16 in murine embryonic stem cells paired with genomic factor localization, nascent transcription, and chromatin accessibility analyses. Over a timecourse of SPT16 depletion, nucleosomes invade loci bound by master pluripotency factors and gene-distal DNaseI hypersensitive sites. Simultaneously, transcription of *Pou5f1* (OCT4), *Sox2, Nanog*, and enhancer RNAs produced at the genes’ associated enhancers are downregulated, suggesting that FACT regulates expression of the pluripotency factors themselves. We find that FACT maintains cellular pluripotency through a precise nucleosome-based regulatory mechanism for appropriate expression of both coding and non-coding transcripts associated with pluripotency.

## Introduction

The process of transcription, or polymerase-driven conversion of a DNA template to RNA, is essential to all life and is highly regulated at all stages (reviewed in (Cramer, 2019; Kornberg & Lorch, 1999; X. Liu, Bushnell, & Kornberg, 2013; Roeder, 2019)). A major barrier to transcription by RNA Polymerase II (RNAPII) is the presence of assembled nucleosomes occluding access to the DNA template (reviewed in (Kujirai & Kurumizaka, 2020; Kwak & Lis, 2013; Lorch & Kornberg, 2020; Lorch & Kornberg, 2017; Venkatesh & Workman, 2015)). A nucleosome consists of a tetramer of two copies each of histones H3 and H4 and two H2A-H2B heterodimers which together form the histone octamer, around which ∼147 base pairs of DNA are wrapped (Lorch & Kornberg, 2020; Luger, Mader, Richmond, Sargent, & Richmond, 1997). Nucleosomes are the basic unit that facilitate DNA compaction into a structure known as chromatin (Lorch & Kornberg, 2020; Luger et al., 1997). Chromatin is highly dynamic and carefully regulated to promote or repress expression of certain genes as dictated by cell signaling, environmental conditions, and master regulators of cell fate. The basic nucleosome can be altered through inclusion of histone variants and histone modifications (reviewed in (Henikoff & Ahmad, 2005; Kouzarides, 2007; Martire & Banaszynski, 2020)). Histone modifications are epigenetic post-translational marks that signify particular regions of chromatin; for example, trimethylation of histone H3 at lysine residue 4 (H3K4me3) is found at regions of active transcription, while acetylation of histone H3 at lysine 27 (H3K27ac) identifies canonical active enhancer marks (reviewed in (Bannister & Kouzarides, 2011; Kouzarides, 2007; Marmorstein & Zhou, 2014)).

In addition to histone variants and histone modifications, chromatin regulation also comes in the form of chromatin-modifying enzymes, including nucleosome remodeling factors that translocate DNA and permit mobilization of nucleosomes to regulate accessibility, and histone chaperones, noncatalytic proteins that are responsible for adding and removing histone components, including both core histones and their variant substitutes (reviewed in (Avvakumov, Nourani, & Cote, 2011; De Koning, Corpet, Haber, & Almouzni, 2007; Hammond, Stromme, Huang, Patel, & Groth, 2017; Ransom, Dennehey, & Tyler, 2010; Venkatesh & Workman, 2015)). To create an RNA product, RNAPII coordinates with these histone chaperones to overcome the physical hindrance of nucleosome-compacted DNA (reviewed in (Formosa, 2012; Hsieh et al., 2013; Kujirai & Kurumizaka, 2020; Kulaeva, Hsieh, Chang, Luse, & Studitsky, 2013; Petesch & Lis, 2012)). RNAPII can facilitate this nucleosome disassembly (Ranjan et al., 2020), but the polymerase is often assisted by the various histone chaperones that can facilitate removal of H2A/H2B dimers (as well as other combinations of histone proteins) and subsequent reassembly after the polymerase has passed (Fei et al., 2018; Lee et al., 2017; Y. Liu et al., 2020; T. Wang et al., 2018). One prominent histone chaperone is the FAcilitates Chromatin Transactions (FACT) complex.

The mammalian FACT complex is a heterodimer composed of a dimer exchange subunit, Suppressor of Ty 16 homolog (SPT16) and an HMG-containing subunit that facilitates localization and DNA binding, Structure-Specific Recognition Protein 1 (SSRP1) (Belotserkovskaya et al., 2003; Y. Liu et al., 2020; G. Orphanides, LeRoy, Chang, Luse, & Reinberg, 1998; G Orphanides, Wu, Lane, Hampsey, & Reinberg, 1999). In *S. cerevisiae*, the system in which much FACT characterization has been done, Spt16 forms a complex with Pob3, assisted by Nhp6, which has been proposed to fulfill the roles of the SSRP1 HMG domain (Brewster, Johnston, & Singer, 1998, 2001; Formosa et al., 2001; G. Orphanides et al., 1998; G Orphanides et al., 1999; Wittmeyer & Formosa, 1997). FACT regulates passage through the nucleosomal roadblock for both RNAPII and replication machinery (Abe et al., 2011; Belotserkovskaya et al., 2003; Belotserkovskaya, Saunders, Lis, & Reinberg, 2004; Formosa, 2008, 2012; Formosa & Winston, 2020; Hsieh et al., 2013; G. Orphanides et al., 1998; G Orphanides et al., 1999; B. C. Tan, Chien, Hirose, & Lee, 2006; Tettey et al., 2019). Given these dual roles in transcription and DNA replication, FACT has been thought to be crucial for cell growth and proliferation (Abe et al., 2011; Belotserkovskaya et al., 2004; Formosa et al., 2001; Garcia et al., 2011; Hertel L., 1999; G Orphanides et al., 1999; B. C. Tan et al., 2006). More recent data has shown that while FACT is not required for cell growth in most healthy adult cell types, FACT is highly involved in cancer-driven cell proliferation as a dependency specific to cancerous cells (Garcia et al., 2013; Kolundzic et al., 2018; Mylonas & Tessarz, 2018; Shen, Formosa, & Tantin, 2018). This dependency has been targeted using a class of FACT inhibitors known as curaxins, with promising results in anticancer drug treatment studies (Chang et al., 2019; Chang et al., 2018; Gasparian et al., 2011). Curaxins inhibit FACT through a trapping mechanism whereby FACT is redistributed away from transcribed regions to other genomic loci, where the complex tightly binds to nucleosomes and cannot be easily removed (Chang et al., 2018). While cancer cell proliferation is FACT-dependent, FACT expression is nearly undetectable in most non-cancerous adult mammalian tissues; indeed, FACT appears to be dispensable for cell viability and growth in most non-cancerous and differentiated cell types (Garcia et al., 2011; Garcia et al., 2013; Safina et al., 2013). Formosa and Winston have recently suggested a unifying model for FACT action wherein cellular FACT dependency results from chromatin disruption and tolerance of DNA packaging defects within the cell (Formosa & Winston, 2020).

While FACT did not initially seem essential for cell proliferation outside of the context of cancer, more recent work has demonstrated heightened FACT expression and novel requirement in undifferentiated (stem) cells (Garcia et al., 2011; Garcia et al., 2013; Kolundzic et al., 2018; Mylonas & Tessarz, 2018; Shen et al., 2018). Stem cell chromatin is highly regulated by well-characterized features, including a largely accessible chromatin landscape and bivalent chromatin, which is epigenetically decorated with both active (e.g., H3K4me3) and repressive (e.g., H3K27me3) modifications (Azuara et al., 2006; Bernstein et al., 2006; de Dieuleveult et al., 2016; Harikumar & Meshorer, 2015; Klein & Hainer, 2020; Meshorer & Misteli, 2006; Vastenhouw & Schier, 2012; Voigt, Tee, & Reinberg, 2013; Young, 2011). Embryonic stem (ES) cells specifically regulate their chromatin to prevent differentiation from occurring until appropriate, thereby preserving their pluripotent state. Pluripotency, or the capacity to mature into most cell types in an adult organism, is maintained by a suite of master regulators that work to repress differentiation-associated genes and maintain expression of genes that promote this pluripotent state, including the well-studied transcription factors OCT4, SOX2, KLF4, MYC, and NANOG, often referred to as master regulators of pluripotency (Chambers et al., 2003; Ding, Xu, Faiola, Ma’ayan, & Wang, 2012; Hall et al., 2009; Kim et al., 2018; Klein & Hainer, 2020; Masui et al., 2007; Mitsui et al., 2003; Pardo et al., 2010; Romito & Cobellis, 2016). While the main functions of these factors are to maintain pluripotency and prevent improper differentiation through regulation of gene expression, a majority of their chromatin binding sites are to gene-distal genomic regions (such as enhancers), suggesting important regulatory functions at these locations (Lodato et al., 2013). These transcription factors, along with chromatin modifiers, form the foundation of gene regulation and provide a molecular basis for pluripotency. FACT has been shown to interact with several pluripotency- and development-associated factors, including OCT4 (Ding et al., 2012; Pardo et al., 2010), WNT (Hossan et al., 2016), and NOTCH (Espanola et al., 2020). Specifically, affinity mass spectrometry has demonstrated an interaction between FACT and OCT4 (Ding et al., 2012; Pardo et al., 2010). In addition, FACT has been functionally implicated in maintaining stem cells in their undifferentiated state (Kolundzic et al., 2018; Mylonas & Tessarz, 2018; Shen et al., 2018). FACT depletion by SSRP1 shRNA knockdown led to a faster differentiation into neuronal precursor cells, along with increased expression of genes involved in neural development and embryogenesis (Mylonas & Tessarz, 2018). In both *C. elegans* and murine embryonic fibroblasts (MEFs), FACT was shown to impede transition between pluripotent and differentiated states; in *C. elegans*, FACT was identified as a barrier to cellular reprogramming of germ cells into neuronal precursors, while in MEFs, FACT inhibition prevented reprogramming to induced pluripotent stem cells (Kolundzic et al., 2018; Shen et al., 2018). These experiments have confirmed a dependency for FACT in pluripotent cells that is not found in differentiated fibroblasts (Kolundzic et al., 2018; Shen et al., 2018). While these data establish FACT as essential in pluripotent cells, the mechanism through which FACT acts within undifferentiated cells to maintain their state is currently unclear. Interestingly, SSRP1 knockout in murine ES cells is viable and shows no effect on expression of the pluripotency factor OCT4 (F. Chen et al., 2020); however, conditional knockout of SSRP1 in mice is lethal due to a loss of progenitor cells resulting in hematopoietic and intestinal failures (Goswami et al., 2022). These disparities may be related to described FACT-independent roles of SSRP1 (Y. Li, Zeng, Landais, & Lu, 2007; Marciano et al., 2018), but nonetheless highlight inconsistencies regarding the role of FACT in pluripotent cells.

Here, we establish a molecular mechanism by which the FACT complex is required for pluripotency in murine ES cells by maintaining the expression of master regulatory transcription factors through their enhancers. As the majority of OCT4, SOX2, and NANOG binding occurs at gene-distal regulatory sites, we sought to determine whether FACT may regulate these factors, along with their regulatory targets, at non-genic locations (Lodato et al., 2013). We identify extensive regulation of non-coding transcription by the FACT complex at cis-regulatory elements such as enhancers and promoters. SPT16 binding is highly enriched at putative enhancers, and transcription of putative enhancer RNAs (eRNAs) is altered between 12 and 24 hours of depletion (2.4% and 34%, respectively), including eRNAs transcribed from enhancers of *Pou5f1, Sox2*, and *Nanog*. Furthermore, we identify co-occupancy between FACT and master regulators of pluripotency and altered nucleosome positioning following a time course depletion of FACT. Together, these data suggest that FACT maintains open chromatin structure at both enhancers and promoters to permit OCT4, SOX2, and NANOG binding and subsequent expression of genes required for pluripotency.

## Results

### Inducible depletion of the FACT complex triggers a reduction in pluripotency

To determine the mechanism through which FACT is critical to stem cell identity, we performed proteasomal degradation of the FACT subunit SPT16 via the auxin-inducible degron (AID) system (Fig. 1A) in murine embryonic stem (ES) cells. Briefly, we used Cas9-directed homologous recombination to insert a mini-AID and 3XV5 tag at the C-terminus of endogenous *Supt16h*, the gene encoding SPT16, in ES cells that have osTIR1 already integrated within the genome (see Methods). Throughout the following described experiments, the osTIR1 cell line, without any AID-tagged proteins, is used as the control cell line (hereafter referred to as “Untagged”). SPT16 protein levels were effectively reduced by proteasomal degradation following 24 hours of treatment with the auxin 3-IAA, and partially reduced after 6 hours, whereas under 6 hours had modest to no reduction in SPT16 levels relative to the vehicle treatment control (EtOH; Fig. 1B, Fig. S1A). We note, as previously established, that depletion of SPT16 triggers a corresponding loss of expression of SSRP1 protein (Fig. S1B) (Safina et al., 2013). Consistent with a role for FACT in stem cell identity and viability, within 24 hours, ES cell colonies began to show phenotypic changes indicative of cellular differentiation, including a loss of alkaline phosphatase activity and morphological changes (Fig. 1C, Fig. S1C). This phenotypic change was most apparent between 24 and 48 hours of FACT depletion; however, most cells could not survive 48 hours of FACT depletion. While it has been suggested that FACT requirement in stem cells is a result of cellular stress induced by trypsinization, we note that cells had been left undisturbed for 48 hours prior to protein depletion, suggesting that trypsinization is unrelated to the differentiation defect or the requirement for FACT (Shen et al., 2018).

**Fig. 1.**
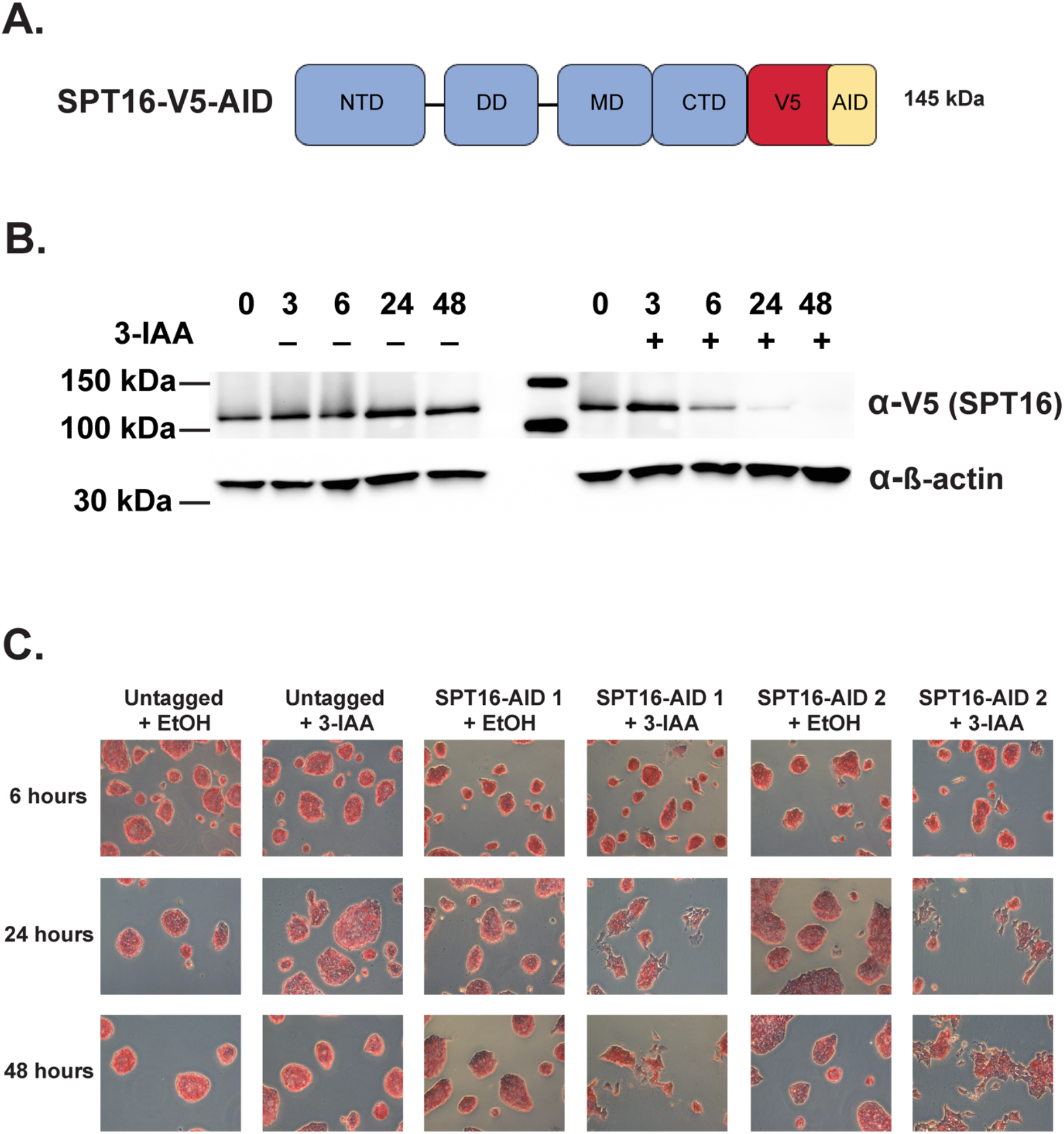
Inducible depletion of SPT16 triggers a loss of pluripotency in ES cells. A. Schematic of auxin-inducible degron (AID) and V5-tagged SPT16 protein. NTD = N-terminal domain, DD = dimerization domain, MD = middle domain, CTD = C-terminal domain, AID = minimal auxin-inducible degron tag, V5 = 3xV5 epitope tag. B. Western blot showing depletion of SPT16 after 0, 3, 6, 24, and 48 treatments with 3-IAA (+) or vehicle control (EtOH, −). 40 µg total protein loaded per lane. Top to bottom, anti-V5 antibody (for tagged SPT16) and anti-β-actin antibody. Representative blot shown from SPT16-V5-AID 1; additional blots can be found in Fig. S1. C. Time course of 3-IAA or EtOH treatment for 6, 24, or 48 hours to deplete SPT16 showing morphological changes following alkaline phosphatase staining. Images are representative of plate-wide morphological changes.

### The FACT complex is enriched at pluripotency factor binding sites

To determine where FACT is acting throughout the genome, we performed the chromatin profiling technique CUT&RUN on the endogenously tagged SPT16-V5 protein (Skene & Henikoff, 2017). Attempts at profiling SPT16 or SSRP1 with antibodies targeting the endogenous proteins were unsuccessful in our hands. SPT16-V5 CUT&RUN recapitulates known FACT binding trends, including a correlation with nascent transcription and with FACT ChIP-seq results (Fig. 2A). However, CUT&RUN also provides heightened sensitivity, allowing for higher resolution profiling and investigation of FACT binding (Hainer, Boskovic, McCannell, Rando, & Fazzio, 2019; Hainer & Fazzio, 2019; Meers, Bryson, Henikoff, & Henikoff, 2019; Skene & Henikoff, 2017). Individual SPT16-V5 CUT&RUN replicates display a higher Pearson correlation than FACT ChIP-seq data, suggesting greater replicability (Fig. S2A).

**Fig. 2.**
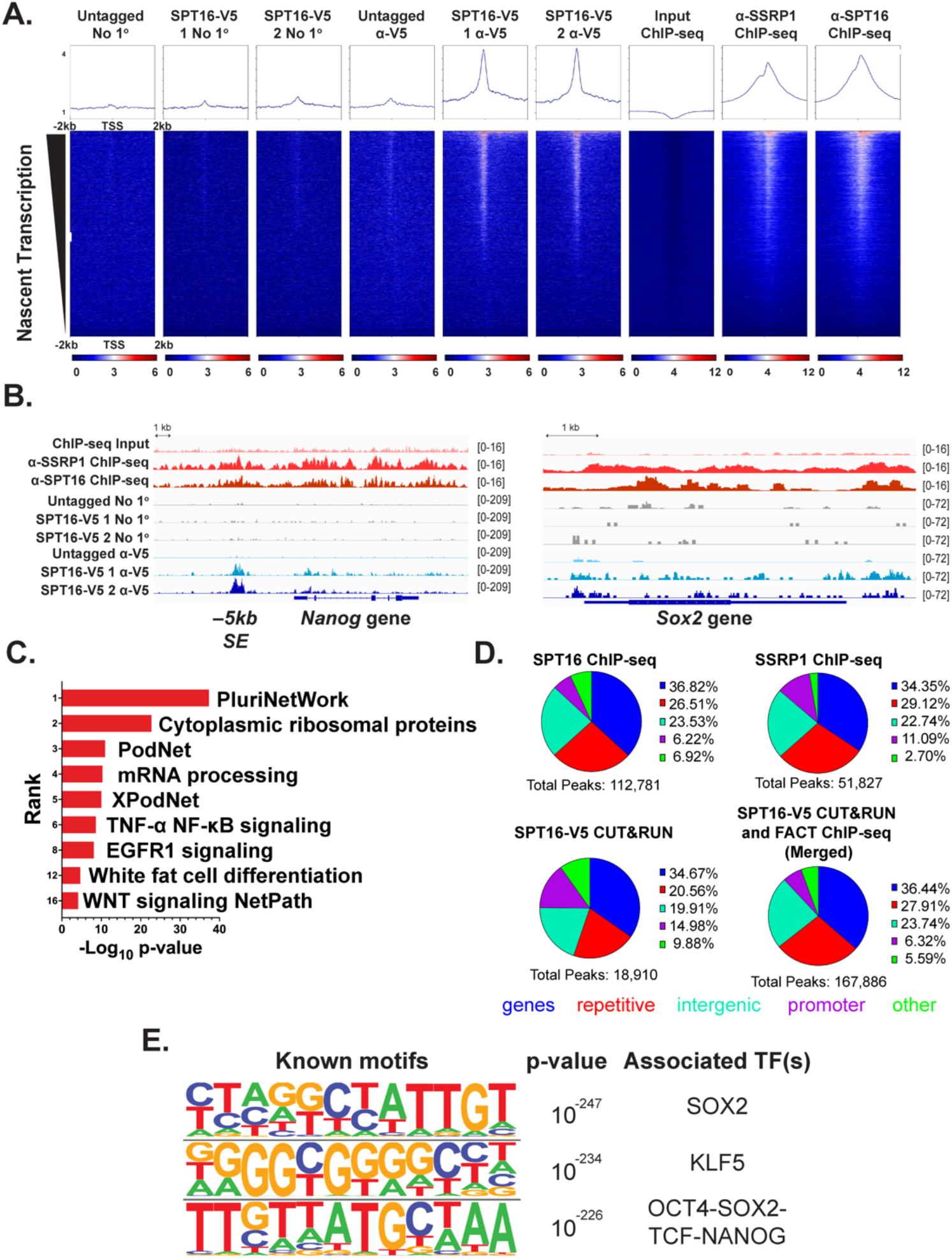
FACT binding is enriched at sites occupied by master regulators of pluripotency. A. SPT16-V5 CUT&RUN and published SPT16 and SSRP1 ChIP-seq data visualized over transcription start sites and sorted by nascent transcription (TT-seq) in control samples (see Fig. 3; ChIP-seq data: GSE90906) (Mylonas & Tessarz, 2018). Averaged replicates are shown as heatmaps +/−2kb from the center of the V5 peak (n = 3 for untagged, n = 2 for each V5-tagged clone, n = 2 for all ChIP-seq experiments). No 1° refers to the negative control experiment where no primary antibody is added, but pA/G-MNase is still added to assess background cutting. B. IGV genome browser track comparing binding trends between CUT&RUN and ChIP-seq data at the *Nanog* (left) and *Sox2* (right) loci. Averaged replicates are shown as a single track (n = 3 for untagged, n = 2 for each V5-tagged clone, n = 2 for all ChIP-seq experiments). C. Pathway analysis of genic SPT16-V5 CUT&RUN peaks identifies enrichment of pluripotency- and differentiation-associated pathways. D. Proportion of peaks called from each dataset corresponding to gene bodies (blue), repetitive regions (red), intergenic regions (mint), promoters (purple, defined as 1 kb upstream of annotated TSSs), and other regions (green). E. The three most significantly enriched sequence motifs of all SPT16-V5 CUT&RUN peaks (n = 4).

To identify and compare FACT-regulated genes from these datasets, we called peaks from CUT&RUN data using SEACR and ChIP-seq data using HOMER (Heinz et al., 2010; Meers, Tenenbaum, & Henikoff, 2019). Overall, FACT ChIP-seq data and SPT16-V5 CUT&RUN data are generally agreeable at peaks called from the orthogonal dataset (Fig. 2A, Fig. S2B-C). In both the SPT16-V5 CUT&RUN data and FACT subunit ChIP-seq, we see strong complex binding at the pluripotency-regulating genes and their distal regulatory elements, such as *Nanog* and *Sox2* (Fig. 2B). We then subjected genic peaks from the CUT&RUN data to Gene Ontology (GO) term analysis, identifying numerous pluripotency- and development-associated pathways (Fig. 2C). Patterns of localization to genomic features were generally similar between experiments (Fig. 2D). We identified 18,910 nonunique peaks called from SPT16-V5 CUT&RUN data, 112,781 nonunique peaks from SSRP1 ChIP-seq data, and 51,827 nonunique peaks from SPT16 ChIP-seq data. CUT&RUN data included more peaks overlapping promoters and unclassified regions, while ChIP-seq peaks contained more repetitive and intergenic regions (Fig. 2D). While we note that more peaks were called from both ChIP-seq datasets, we caution against interpreting raw peak numbers due to greatly differing sequencing depth and false discovery rates employed by the respective peak-calling algorithms.

To assess the association between FACT and pluripotency orthogonally, we performed sequence motif analysis of all CUT&RUN peaks using HOMER (Fig. 2E) (Heinz et al., 2010). The top three most enriched sequence motifs were those recognized by the transcription factors SOX2, KLF5, and OCT4-SOX2-TCF-NANOG, all of which regulate cellular pluripotency or differentiation (Bourillot & Savatier, 2010; Chambers et al., 2003; Hall et al., 2009; Klein & Hainer, 2020; Masui et al., 2007; Mitsui et al., 2003; Pardo et al., 2010). Together, these results led us to propose that FACT may maintain pluripotency of ES cells through coordinated co-regulation of target genes with the master regulators of pluripotency.

### FACT regulates expression of the master regulators of pluripotency

While we identified FACT occupancy over pluripotency genes, it remained unclear whether FACT directly regulates the expression of the master regulators of pluripotency themselves. We therefore performed nascent RNA sequencing (TT-seq) following depletion of SPT16 for a direct readout of FACT’s effects on transcription of these regulators. To assess the effects of SPT16 depletion on transcription over time, we performed a time course of 3, 6, 12, and 24 hours of IAA treatment. Consistent with our analysis of protein depletion (Fig. 1B), we identified few differentially transcribed genes prior to morphological indicators of cellular differentiation (within 6 hours; Table 1). We performed RT-qPCR at 3 and 6 hours of depletion (Fig. S3D-F) and confirmed that *Supt16, Ssrp1, Pou5f1, Sox2*, or *Nanog* transcript abundance had not changed, suggesting that moderate levels of FACT protein are sufficient to sustain pluripotency (Fig. S3D-F). After 12 hours of depletion, however, cells begin to differentiate, and pluripotency factor expression declines (Figs. 3A-B, S3A-B). As pluripotency factor expression does not decline prior to complete depletion of SPT16, we infer that FACT expression is required to maintain pluripotency, potentially by regulating expression of these important transcription factors.

**Table 1.** Significantly altered mRNAs, PROMPTs, and DHS-associated ncRNAs at 0, 3, 6, 12, and 24h of SPT16 depletion. Control samples and SPT16-depleted samples were pooled between cell lines for downstream analyses. Only transcripts with an adjusted p-value of < 0.05 are displayed (analyzed with DESeq2).

| Hours Depleted | 0 | 3 | 6 | 12 | 24 |
| --- | --- | --- | --- | --- | --- |
| Genes Up | 3 | 58 | 214 | 1366 | 5398 |
| Genes Down | 53 | 5 | 174 | 1932 | 5000 |
| PROMPTs Up | 0 | 4 | 4 | 95 | 2984 |
| PROMPTs Down | 0 | 0 | 0 | 59 | 1831 |
| ncRNAs Up | 0 | 4 | 38 | 203 | 8743 |
| ncRNAs Down | 0 | 0 | 34 | 323 | 5789 |

**Fig. 3.**
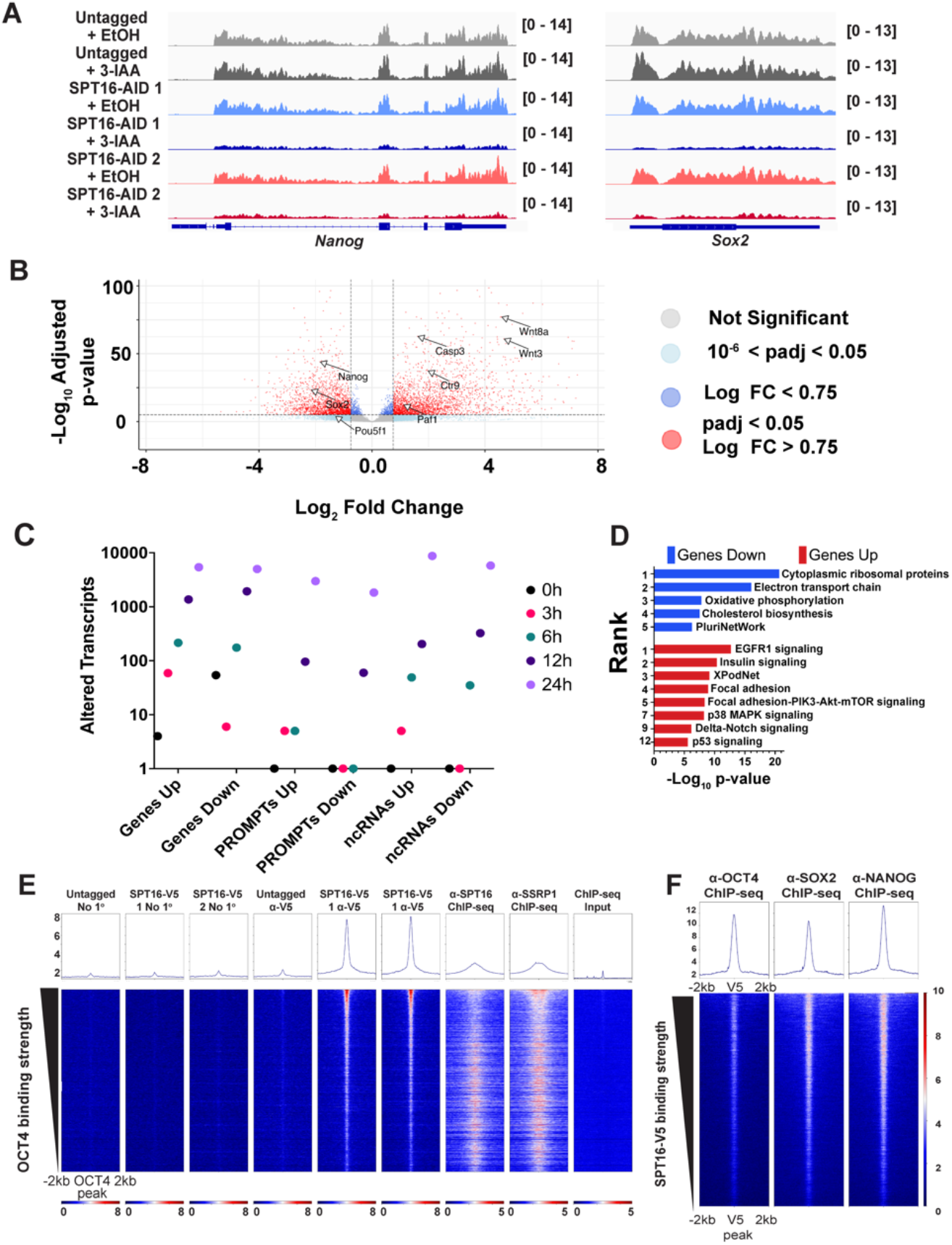
Depletion of FACT disrupts transcription of master regulators of pluripotency. A. IGV genome browser tracks showing nascent transcription from TT-seq experiments over the *Nanog* (left) and *Sox2* (right) genes following 24-hour 3-IAA treatment to deplete SPT16. Averaged replicates are shown as a single track, oriented to the genic strand (n = 3) B. Volcano plot of differential gene expression after 24 hours of treatment (analyzed with DESeq2). Red points indicate significant changes (padj < 0.05, log_2_ fold change > 0.75). Light blue points are significant changes by p-value but below the fold change cutoff, while dark blue points are significant changes by log_2_ fold change but below the p-value cutoff. C. Dot plot depicting the number of differentially expressed genes, PROMPTs, and ncRNAs transcribed from gene-distal DNaseI hypersensitive sites (DHSs) (DNase-seq from GSM1014154) (Consortium, 2012; Davis et al., 2018; Thurman et al., 2012) (Analyzed with DESeq2). Number of transcripts in each category are provided in Table 1. One count was added to each category for plotting. D. Pathway analysis of differentially expressed genes following 24-hour 3-IAA treatment to deplete SPT16. Y-axis indicates Wikipathways enrichment ranking. E. SPT16-V5 CUT&RUN binding enrichment over gene-distal OCT4 ChIP-seq peaks (ChIP-seq from GSE11724) (Marson et al., 2008). Merged replicates are shown as heatmaps +/−2kb from the center of the OCT4 ChIP-seq peak (n = 3 for untagged, n = 2 for each V5-tagged clone, n = 2 for all ChIP-seq experiments; ChIP-seq from GSE11724) (Marson et al., 2008). F. OCT4, SOX2, and NANOG enrichment over SPT16-V5 CUT&RUN peaks. Averaged replicates shown (n = 1 for OCT4, n = 2 for SOX2 and NANOG; ChIP-seq from GSE11724) (Marson et al., 2008).

Intriguingly, partial depletion of SPT16 (≤6 hours) is accompanied by more upregulation of transcription than decreased transcription (Fig. 3C, S4, Table 1); concordant with modest visible differentiation beginning (Fig. 1C), however, we identified more reduced transcription (1,932, 11%) than upregulation (1,366, 7.6%) of protein coding genes at 12 hours of depletion (Fig. 3C, S4, Table 1). At 24 hours, slightly more genes were upregulated (5,398, 27%) than reduced (5,000, 25%) (Fig. 3B-C, S4, Table 1). Of the genes encoding master pluripotency factors, only *Nanog* was significantly reduced within 12 hours of treatment (Fig. 3A, S3A-B), while all four Yamanaka factors (*Pou5f1, Sox2, Klf4*, and *Myc*) and *Nanog* were significantly reduced after 24 hours (Fig. 3A, S3A-B). At 24 hours of depletion, transcription elongation factors were significantly upregulated, such as subunits of the Polymerase-Associated Factors (PAF1) complex, the DRB Sensitivity Inducing Factor (DSIF) member SPT4A, and the histone chaperone SPT6 (Fig. 3B, Fig. S3C). SPT6 has been shown to maintain ES cell pluripotency through Polycomb opposition and regulation of superenhancers (A. H. Wang et al., 2017). Heightened expression of transcription elongation factors may be the result of a compensatory mechanism through which FACT-depleted cells attempt to overcome this deficiency or the result of direct repression of these factors by FACT.

To identify cellular processes critically regulated by FACT, we subjected differentially transcribed genes to pathway analysis after 24 hours of treatment and identified enrichment for the pluripotency network among genes with reduced transcription, while numerous signaling pathways were enriched among the genes with increased transcription (Fig. 3D). As OCT4, SOX2, and NANOG protein expression levels are maintained for 3-5 days in ES cells deprived of LIF (Ee et al., 2017), we infer a direct dependency of the master regulators on FACT; upon FACT depletion, ES cells are forced to differentiate by inability to maintain OCT4, SOX2, and NANOG expression. Given the extensive connections between FACT and pluripotency regulators, we next sought to further characterize this regulatory dynamic.

### FACT co-occupies gene distal regions bound by OCT4, SOX2, and NANOG

Having established that FACT regulates expression of the important pluripotency-regulating genes, we attempted to identify whether this regulation occurs at the genes themselves or at distal regulatory elements. As a majority of OCT4, SOX2, and NANOG binding sites are gene-distal (X. Chen et al., 2008; Lodato et al., 2013) and previously published FACT subunit ChIP-seq correlates poorly with genes that change expression upon SSRP1 knockdown (Mylonas & Tessarz, 2018), we hypothesized that FACT may also bind at gene-distal regulatory sites, especially those sites bound by OCT4, SOX2, and NANOG.

Indeed, both SPT16-V5 CUT&RUN and previously published FACT subunit ChIP-seq (Mylonas & Tessarz, 2018) show strong occupancy over gene-distal OCT4 ChIP-seq peaks, suggesting co-regulation of pluripotency factor targets (Fig. 3E). Orthogonally, we analyzed published OCT4, SOX2, and NANOG ChIP-seq data (Marson et al., 2008) and visualized over SPT16-V5 CUT&RUN peaks (Fig. 3F). All three pluripotency factors display enriched binding at SPT16-V5 binding sites, supporting the idea of co-regulation by FACT and pluripotency factors.

Finally, as there is a known interaction between OCT4 and acetylation of histone H3 at lysine 56 (H3K56ac) (Y. Tan, Xue, Song, & Grunstein, 2013; Xie et al., 2009), we hypothesized that FACT binding may correlate with H3K56ac. In support of this hypothesis, FACT and H3K56ac are known to interact in *S. cerevisiae* (McCullough et al., 2019). As such, we examined whether this interaction is conserved in ES cells. We plotted SPT16-V5 CUT&RUN and published FACT subunit ChIP-seq data over published H3K56ac ChIP-seq peaks (Fig. S2D; H3K56ac ChIP-seq: GSE47387 (Y. Tan et al., 2013)). While FACT does not appear enriched directly over H3K56ac peaks, FACT is highly enriched immediately flanking the H3K56ac peaks. The association between FACT and H3K56ac further highlights FACT’s role in pluripotency maintenance, given the previously established interplay between OCT4 and H3K56ac. We caution, however, against overinterpreting this trend, due to poor specificity of the H3K56ac antibody and low abundance (<1% of total H3 loci) in mammalian cells (Pal et al., 2016). Together, our analyses show that FACT co-localizes at pluripotency-associated sites, including gene-distal regulatory elements.

### SPT16 depletion alters non-coding transcription at gene-distal regulatory sites

FACT binding is strongly enriched at many promoters of genes displaying expression changes following FACT depletion but not at unchanged genes, yet there are other promoters of genes with altered expression following FACT depletion where FACT binding is not detected (Fig. S3A-F); as such, FACT may maintain or repress expression of these target genes through gene-distal regulatory elements. As gene-distal DHSs are often sites of non-protein-coding transcription, including enhancers where enhancer RNAs (eRNAs) are produced, we sought to determine whether FACT localization to gene distal OCT4, SOX2, and NANOG bound putative enhancers may regulate non-coding transcription known to arise from these regions (reviewed in (Kaikkonen & Adelman, 2018; W. Li, Notani, & Rosenfeld, 2016; Patty & Hainer, 2020)). Using our time course depletion of SPT16 followed by TT-seq, we identified FACT-dependent transcription of eRNAs from known superenhancers of the *Pou5f1, Sox2*, and *Nanog* genes (Fig. 4A, Fig. S5). Out of 70,586 putative regulatory regions (defined as gene-distal DNaseI hypersensitive sites), 57,954 were sites of nascent transcription detected in our TT-seq datasets, the majority of which are likely to encode eRNAs (Table 1, Fig. 3C, S4). In analyzing our 24-hour TT-seq data after FACT depletion, we identified 14,532 FACT-regulated ncRNAs (26%), with more ncRNAs derepressed (15%, 8,743) than repressed (11%, 5,789) by FACT depletion (Table 1, Fig. 3C, 4B, S4). Assuming that each ncRNA is paired with (and potentially regulates) its nearest gene, we performed pathway analysis on putative ncRNA regulatory targets (Fig. 4C). Among the most significantly enriched categories for putative targets of upregulated ncRNAs were mechanisms associated with pluripotency, white fat cell differentiation, and WNT signaling, while putative targets of downregulated ncRNAs were enriched for pluripotency networks, TGF-ß signaling, and WNT signaling (Fig. 4C).

**Fig. 4.**
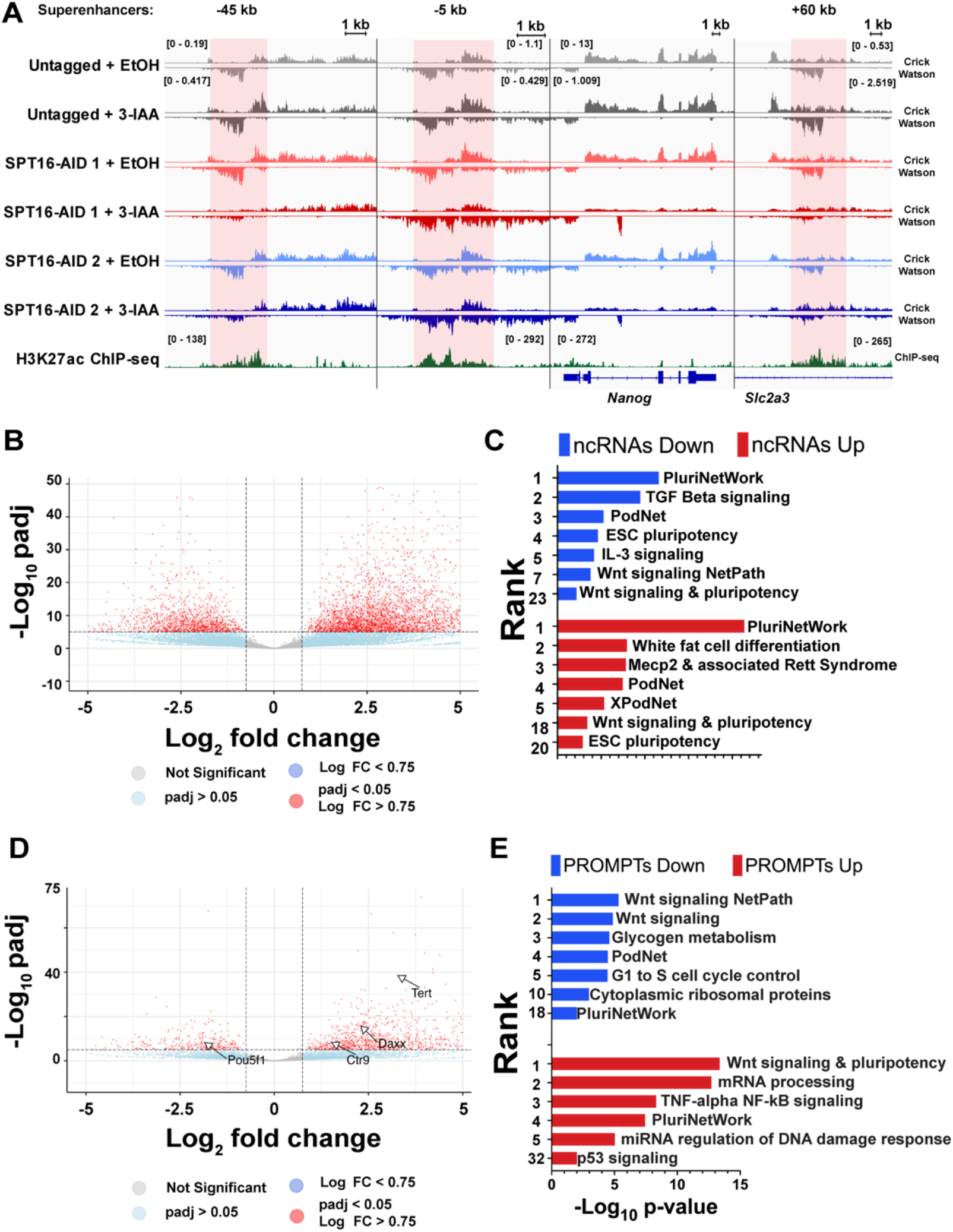
Transient transcriptome sequencing identifies FACT-dependent regulation of non-coding RNAs. A. IGV genome browser tracks showing nascent transcription (TT-seq) over the *Nanog* gene and three *Nanog* superenhancers following 24-hour 3-IAA treatment to deplete SPT16, along with published H3K27ac ChIP-seq data. Three individually scaled windows are shown to highlight eRNA transcription from the superenhancers (shaded red) and *Nanog* gene (shaded blue). Merged replicates are shown as a single track (n = 3, n = 1 for H3K27ac ChIP-seq; ChIP-seq from GSE32218) (Consortium, 2012; Davis et al., 2018; Thurman et al., 2012). B. Volcano plot of differential non-coding RNA expression (analyzed with DESeq2). Red points are significantly changed ncRNAs (padj < 0.05, log_2_ fold change > 0.75). Dark blue points are significantly changed by p-value but below the fold change cutoff, while light blue points are significantly changed by log_2_ fold change but below the p-value cutoff. Labeled arrows denote closest genes to the indicated ncRNA. C. Pathway analysis of differentially expressed ncRNAs following 24-hour 3-IAA treatment to deplete SPT16. Y-axis indicates pathway enrichment ranking. D. As in 4B, but for differentially expressed PROMPTs. E. As in 4C, but for PROMPTs.

Taking only the ncRNAs transcribed from regions marked by both a DHS and either H3K4me1 or H3K27ac as putative eRNAs, we identified 11,964 transcripts, with 18% of putative eRNAs derepressed (2,701) and 16% downregulated (2,439) upon FACT depletion (Table 1, Fig. 3C, S4). Because a majority of OCT4 binding sites are gene-distal, and because FACT binds at gene-distal DHSs and gene-distal OCT4 binding sites (Figs. 3E-F), we sought to determine whether FACT regulates these ncRNAs as a possible means of pluripotency maintenance. Therefore, to examine trends at well-defined enhancers of pluripotency factors, we determined nascent transcription from previously annotated superenhancers known to be marked by eRNA transcription (Blinka, Reimer, Pulakanti, & Rao, 2016; Y. Li et al., 2014; Whyte et al., 2013) (Fig. 4A, Fig. S5A-B).

We next sought to identify putative regulation by FACT of genes via proximal regulatory elements—specifically promoter upstream transcripts (PROMPTs; also referred to as upstream antisense noncoding RNAs or uaRNAs). PROMPTs were identified by genomic location (within 1 kb of an annotated TSS and transcribed divergently to the mRNA); 4,815 PROMPTs were significantly altered by FACT depletion out of 23,256 expressed putative PROMPTs (padj < 0.05; Fig. 4D). More PROMPTs were repressed by FACT than stimulated, with 13% significantly increasing (2,984) and 7.9% significantly decreasing (1,831) (Fig. 3C, 4D, S4, Table 1). Regulation of approximately 20% of putative PROMPTs remains in line with known roles for transcriptional regulation by FACT, and repression of PROMPTs is consistent with FACT’s known role in preventing cryptic transcription *S. cerevisiae* (C. Jeronimo, Watanabe, Kaplan, Peterson, & Robert, 2015; Mason & Struhl, 2003). We subjected identified PROMPTs to pathway analysis by assignment to the nearest gene as in Fig. 4C and identified pluripotency-and differentiation-associated pathways among the most enriched pathways (Fig. 4E). Among the most affected classes of FACT-regulated genes are those that regulate pluripotency and stem cell identity (Fig. 3A-F). Expression of these pluripotency factors is regulated by enhancers and superenhancers; as eRNA transcription from these gene-distal regulatory regions is compromised following FACT depletion (Fig. 4A, Fig. S5A-B), the mechanism through which FACT regulates stem cell pluripotency appears to depend on these enhancers.

### FACT binds to gene-distal putative enhancers and not putative silencers

Based on our finding that putative eRNAs require FACT for appropriate expression, we assessed FACT binding at a number of features defining regulatory regions, including H3K27ac ChIP-seq peaks (Fig. 5A), H3K4me1 ChIP-seq peaks (Fig. 5B), gene-distal DHSs (Fig. 5C), and H3K56ac ChIP-seq sites (Fig. 5D). At each of these sites marking putative regulatory regions (typically enhancers), FACT is bound according to both SPT16-V5 CUT&RUN data and FACT subunit ChIP-seq data (Mylonas & Tessarz, 2018). To confirm that FACT is present at putative enhancers, we defined DHSs that were also decorated by either H3K27ac or H3K4me1, two putative enhancer marks, and visualized FACT localization profiling at these sites (Fig. 5E). Indeed, both CUT&RUN and previously published ChIP-seq showed enrichment of FACT binding at putative enhancers. Although FACT binds many regulatory regions marked by DHSs, we note that FACT binding is not enriched at putative silencers, defined by the presence of a TSS-distal DHS and an H3K27me3 ChIP-seq peak (Fig. S7A-D). To determine whether FACT depletion may stimulate transcription from all regulatory elements marked by DHSs, we examined FACT-dependent transcription from these putative silencers. FACT does not appear to stimulate transcription from putative silencers, as there is no discernable enrichment for FACT binding, nor is there an increase in transcription from these regions following FACT depletion (Fig. S7A-D).

**Fig. 5.**
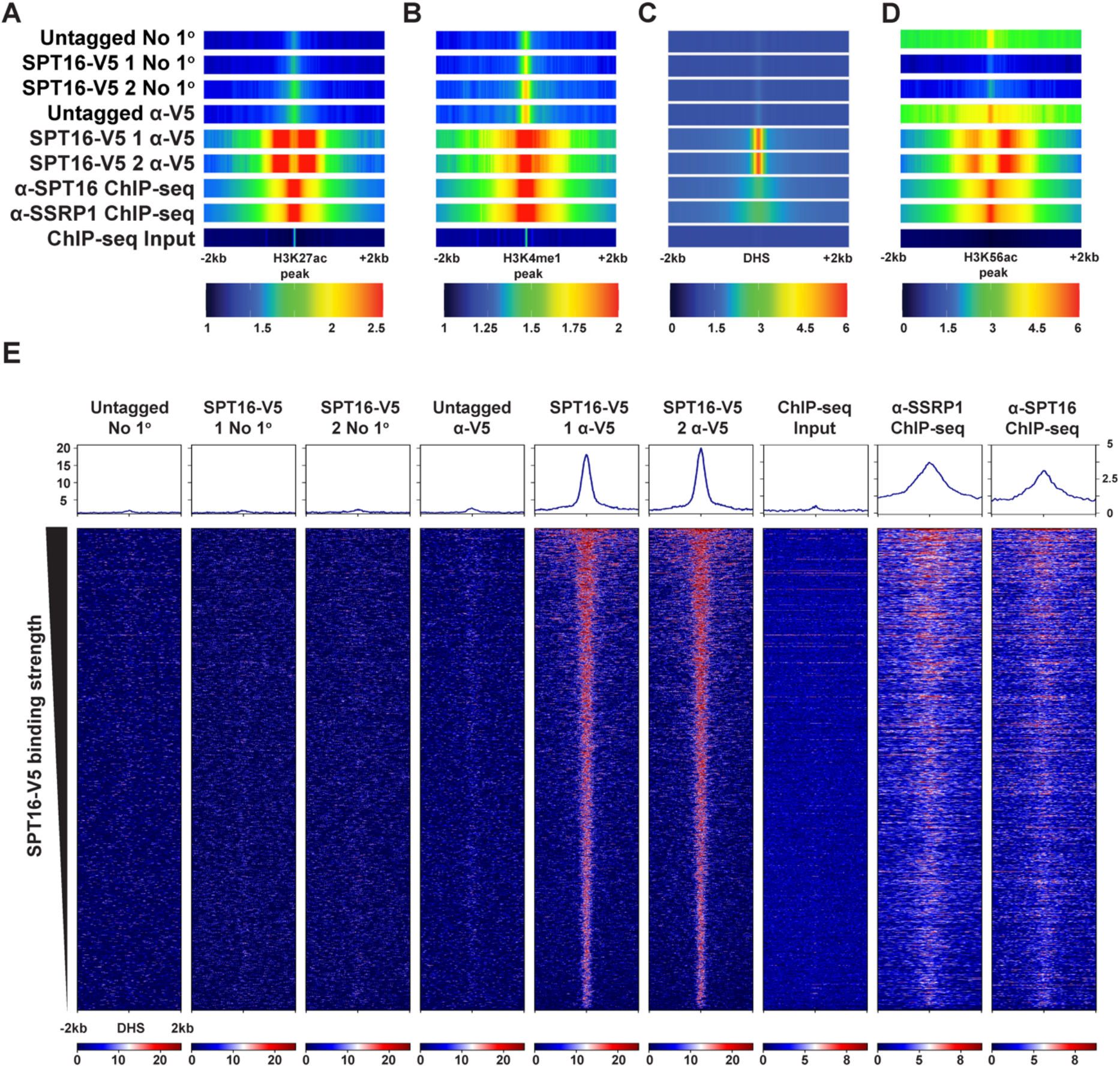
FACT binds to putative gene-distal regulatory regions genome-wide. A-D. SPT16-V5 CUT&RUN, SPT16 ChIP-seq (GSE90906), and SSRP1 ChIP-seq (GSE90906) data visualized as one-dimensional heatmaps (Mylonas & Tessarz, 2018). Each row represents the average of technical replicates, while biological replicates are displayed separately (n = 3 for untagged, n = 2 for each V5-tagged clone, n = 2 for all ChIP-seq experiments). Visualized at A. H3K27ac ChIP-seq peaks +/−2kb (GSE32218) (Consortium, 2012; Davis et al., 2018; Thurman et al., 2012). B. H3K4me1 ChIP-seq peaks +/−2kb (GSE31039) (Consortium, 2012; Davis et al., 2018; Thurman et al., 2012). C. Gene-distal DNaseI hypersensitive sites +/−2kb (GSM1014154) (Consortium, 2012; Davis et al., 2018; Thurman et al., 2012). D. Gene-distal H3K56ac peaks overlapping nonunique peaks called from SPT16-V5 CUT&RUN, SPT16 ChIP-seq, and SSRP1 ChIP-seq (GSE90906) (Mylonas & Tessarz, 2018). E. SPT16-V5 CUT&RUN, SPT16 ChIP-seq, and SSRP1 ChIP-seq data visualized at SPT16-V5-bound putative enhancers, defined as DHSs (GSM1014154) overlapping H3K4me1 or H3K27ac ChIP-seq peaks (GSE32218 and GSE31039) +/− 2kb (Consortium, 2012; Davis et al., 2018; Thurman et al., 2012). F. Metaplots of ATAC-seq data showing differential chromatin accessibility following 24 hours of 3-IAA treatment to deplete SPT16 vs. vehicle, visualized at FACT-bound putative enhancers (as defined in 5D) +/− 2kb. Biological replicates are displayed separately in red and blue (n = 1), while untagged samples are shown in gray (averaged; n = 2). Standard error is shaded in either direction.

To summarize the findings thus far, FACT displays both repressive and permissive effects on transcription arising from genes and gene-distal regulatory regions (Fig. 3B-C, Fig. 4B-E). While FACT stimulates and impedes transcription through direct action at some promoters, a large class of genes with FACT-regulated transcription are not bound by FACT, suggesting gene-distal regulatory mechanisms (Fig. S6A-F). Given the overlap between FACT binding and various enhancer-associated histone modifications (H3K27ac, H3K4me1, H3K56ac; Fig. 5A-D), gene-distal regulation may occur predominantly through association with enhancers of FACT-regulated genes.

### SPT16 depletion results in decreased chromatin accessibility over FACT-bound sites

As FACT is a histone chaperone that can exchange histone H2A/H2B dimers, we hypothesized that FACT may maintain pluripotency by enforcing appropriate chromatin accessibility, including at the gene-distal sites where OCT4, SOX2, and NANOG bind. To identify changes in chromatin accessibility upon FACT depletion, we performed ATAC-seq across a 3h, 6h, 12h, and 24h time course of IAA treatment. Consistent with the localization trends described in Figs. 3E-F and 5, FACT depletion leads to reduced accessibility directly over gene-distal DHSs after 6 hours (Fig. 6A). Specifically, at both 12 and 24 hours there is lower chromatin accessibility in SPT16-depleted cells relative to vehicle-treated or untagged controls. Unsurprisingly, we see this same trend when visualized over SPT16-V5 peaks, many of which overlap these gene-distal DHSs (Fig. 6B). We identified a similar trend over genes (Fig. 6C), perhaps due to the loss of FACT’s established role in replacing histones in the wake of RNAPII at highly transcribed genes, or the inability of FACT to facilitate pause release from more lowly transcribed genes (Farnung, Ochmann, Engeholm, & Cramer, 2021; Tettey et al., 2019). In light of FACT’s consistent role in preserving accessibility at both genes and gene-distal regulatory elements, we reexamined our FACT localization data at these regions. We identified similar trends of FACT binding at both gene-distal and genic SPT16-V5 binding sites, implying similar regulation of both categories (Fig. S5A). Furthermore, we identified little distinction between promoter- and distal accessibility changes upon FACT depletion. We see a marked decrease in chromatin accessibility directly over the DHS, indicating that FACT is necessary to maintain accessible chromatin at putative enhancers (Fig. 6D). Together, these data suggest a mechanism of nucleosome-filling, wherein FACT typically assists in maintaining accessible chromatin at gene-distal regulatory elements.

**Fig. 6.**
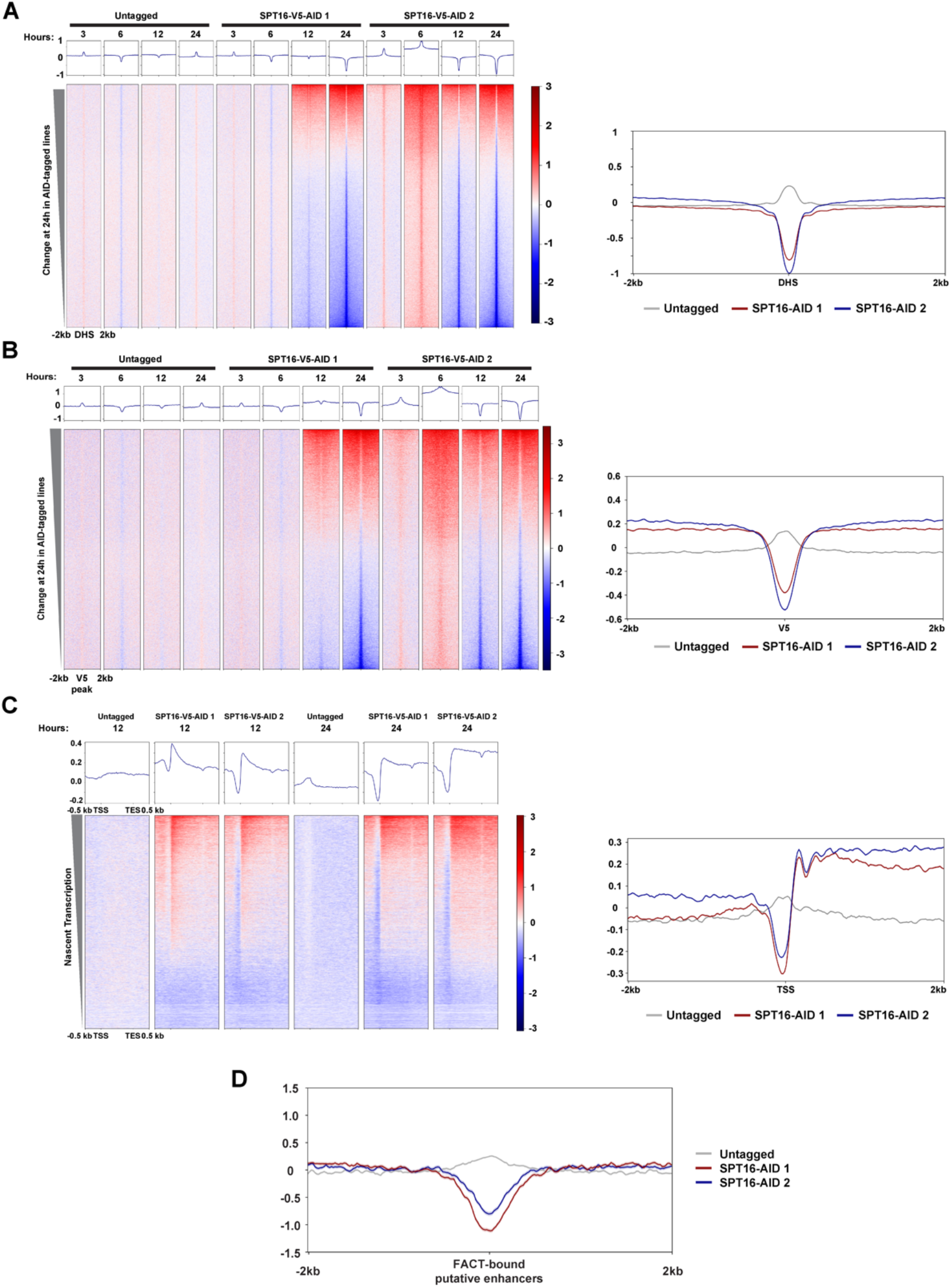
FACT depletion has distinct effects on chromatin accessibility at SPT16-V5 binding sites and gene regulatory regions. A. Differential chromatin accessibility visualized over gene-distal DHSs, +/−2kb, at 3, 6, 12, and 24 hours of treatment. Higher signal indicates more accessible chromatin in 3-IAA-treated samples than in EtOH-treated samples at the indicated timepoint, with the exception of untagged samples (3-IAA:3-IAA ratio) (n = 1 per timepoint). Metaplot of data from 24 hours shown at right. B. As in A, but visualized over SPT16-V5 binding sites identified in Fig. 2. C. Metagene plots depicting changes in chromatin accessibility over FACT-bound gene bodies after 12-24 hours of FACT depletion. Data are sorted by nascent transcription in control samples as in Fig. 2A. Differential signal was calculated as in A. D. Metaplot depicting change in chromatin accessibility at 24 hours of treatment over putative enhancer regions as defined in Fig. 5.

### SPT16 depletion leads to increased nucleosome occupancy over FACT-bound locations

For a more precise understanding of changes to nucleosome occupancy and positioning, we performed micrococcal nuclease digestion followed by deep sequencing (MNase-seq) following FACT depletion after 24 hours of 3-IAA treatment. MNase-seq results suggest a consistent mechanism of nucleosome-filling at FACT-bound regulatory regions genome-wide (Fig 7). Visualizing MNase-seq data at bound peaks called from SPT16-V5 CUT&RUN data, we observe an overall increase in nucleosome occupancy directly over SPT16-V5 peaks following SPT16 depletion (Fig. 7A). Consistent with FACT binding trends identified in Fig. 2, this mechanism of nucleosome filling is not restrained to genic FACT-binding sites; at gene-distal DNaseI hypersensitive sites (DHSs), used as a proxy for gene-distal regulatory regions, a similar phenomenon of nucleosome filling occurs (Fig. 7B, S8). At OCT4, SOX2, and NANOG binding sites, we also observe an increase in nucleosome occupancy after FACT depletion (Fig. 7C-E). In examining annotated TSSs, we also observed increased nucleosome occupancy directly over promoter regions and altered occupancy of downstream genic nucleosomes (Fig. 7F), in agreement with our ATAC-seq data (Fig. 6C). Together with the ATAC-seq data, these data demonstrate increased nucleosome occupancy upon SPT16 depletion. We suggest a model (Fig. 8) where FACT maintains pluripotency through both gene-proximal and gene-distal regulation of pluripotency transcription factors.

**Fig. 7.**
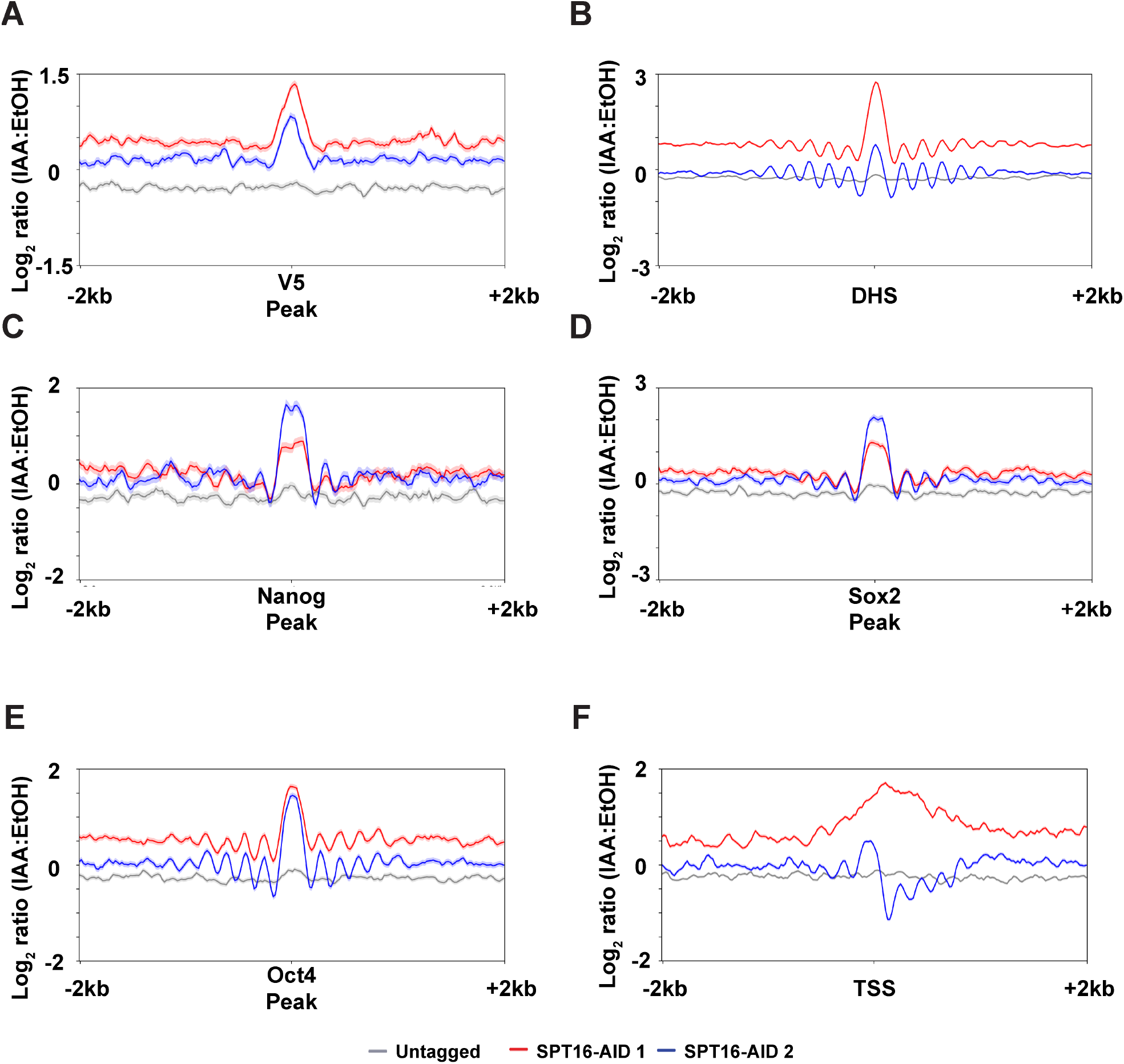
SPT16 depletion disrupts nucleosome positioning at pluripotency-associated sites. A-F. Metaplot differential nucleosome occupancy between 3-IAA and vehicle-treated samples. Averaged replicates are shown as a single line (n = 3 for untagged, n = 2 for each AID-tagged clone). Tagged samples are shown in red and blue, while untagged samples are shown in grey. Shaded area indicates standard error. Visualized over A. peaks called from SPT16-V5 CUT&RUN +/−2kb. B. gene-distal DNaseI hypersensitive sites (DHSs) +/−2kb (DNase-seq from GSM1014154) (Consortium, 2012; Davis et al., 2018; Thurman et al., 2012) C. NANOG ChIP-seq binding sites, +/−2kb (ChIP-seq from GSE11724) (Marson et al., 2008) D. SOX2 ChIP-seq binding sites, +/−2kb (ChIP-seq from GSE11724) (Marson et al., 2008) E. OCT4 ChIP-seq binding sites, +/−2kb (ChIP-seq from GSE11724) (Marson et al., 2008) F. TSSs, +/−2kb.

**Fig. 8.**
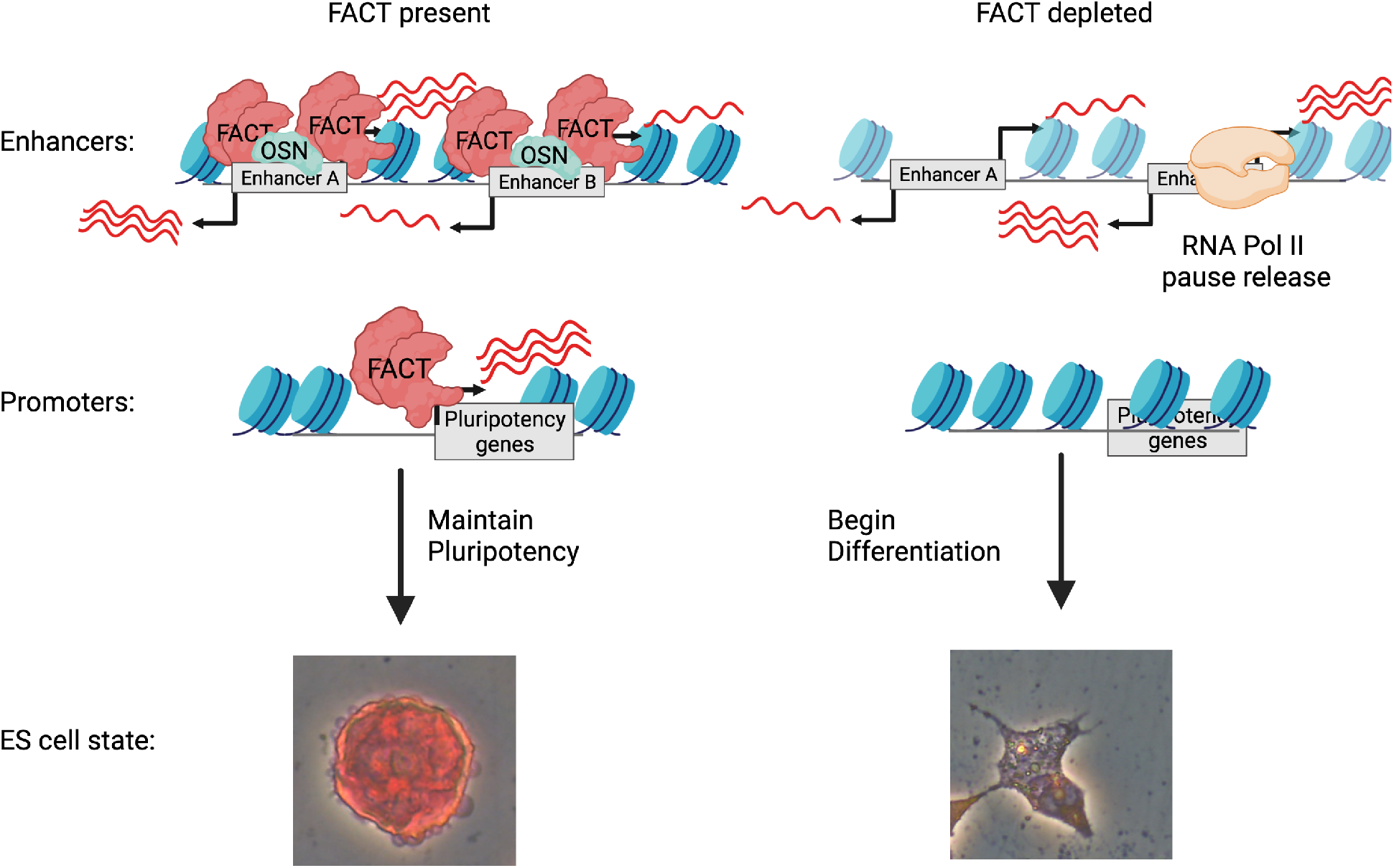
FACT maintains ES cell pluripotency through regulation of pluripotency factor expression. FACT binds to gene distal cis-regulatory elements (enhancers) and regulates both ncRNA transcription and nucleosome occupancy at these regulatory locations to permit appropriate expression of mRNAs. FACT may regulate expression of both coding and non-coding transcripts through maintenance of RNAPII pausing. When FACT is depleted through 3-IAA treatment, transcription and nucleosome occupancy at cis-regulatory elements is disrupted and mRNA expression is altered. These changes result in a loss of pluripotency and ES cells differentiate. OSN = OCT4, SOX2, and NANOG. Image was created with Biorender.com.

## Discussion

### FACT is an essential regulator of stem cell pluripotency

The role for FACT in pluripotent cells has drawn recent interest but remained mechanistically unclear. Here we provide an analysis of FACT function in murine embryonic stem (ES) cells. Our data indicate that FACT regulates pluripotency factors through maintenance of master pluripotency regulators themselves and through gene-distal mechanisms. Given the genomic loci at which FACT binds and the effects of FACT depletion on their transcription, FACT likely performs dual roles in transcriptional regulation: facilitation of pluripotency through both coding and non-coding pluripotency-promoting elements, and repression of differentiation-promoting elements. Based on these data, we propose a model where FACT maintains paused RNAPII at transcribed regions to repress transcription of differentiation-associated genes and non-coding RNAs that may themselves repress pluripotency factors (Fig. 8). Simultaneously, FACT maintains expression of pluripotency factors, through both genic (RNAPII pause release) and gene-distal (enhancer-driven) mechanisms. FACT tends to repress transcription of both coding and non-coding elements at approximately 1.5 times the amount the complex stimulates transcription of coding-and non-coding elements, based on number of differentially regulated transcripts called by DESeq2 data (Fig. 3B-C, 4B, 4D, S4, Table 1). The amount of FACT-dependent mRNA transcription (both stimulated and repressed) are largely consistent between our data and experiments performed in *S. cerevisiae* (Feng et al., 2016), ES cell lines (F. Chen et al., 2020), and in a mouse model (Goswami et al., 2022), suggesting conservation of FACT function throughout eukaryotes.

### FACT regulates chromatin accessibility and transcription at gene-distal regulatory sites

Elucidating a mechanism of FACT action remains complicated by the duality of the complex’s roles; at some loci, FACT works to repress transcription of regulatory elements, while others are positively regulated to promote transcription of their genic targets (Fig. 4B, 4D). Indeed, FACT’s role at gene-distal regulatory elements seems to mirror the complex’s role at genic regions, facilitating removal of nucleosomes to maintain expression when necessary, and reconstruction of nucleosomes to limit expression. While our data indicate that FACT’s more prominent role at gene-distal DHSs is repression of transcription, the complex both facilitates and impedes coding and non-coding transcription, through direct (and likely some indirect) mechanisms (Fig. S2B). The classes of RNAs regulated by FACT do not appear solely categorized by ES cell requirement, however, as GO-term analysis identified many distinct pathways among the most enriched for each class of RNA (Fig. 4C, E). Given the extensive non-coding transcription that arises from gene-distal regulatory elements (Patty & Hainer, 2020), the act of transcription by RNAPII may be the driving force behind increased chromatin accessibility at transcribed regions upon FACT depletion. Whereas FACT would typically reset the nucleosome array in the wake of RNAPII, transcription in FACT-depleted cells appears to stimulate chromatin accessibility and further transcription, in line with prior suggestions (Farnung et al., 2021; Formosa & Winston, 2020).

### Ideas and Speculation

It is tempting to speculate that FACT must maintain accessible chromatin for interaction by the master regulators of pluripotency themselves; however, established pioneering activity by OCT4 and SOX2 suggests that the master regulators are not entirely dependent on FACT action (Dodonova, Zhu, Dienemann, Taipale, & Cramer, 2020; Michael et al., 2020; Soufi et al., 2015; C. Tan & Takada, 2020). FACT depletion has been shown to redistribute histone marks in *D. melanogaster* and *S. cerevisiae*; therefore, disruption of pluripotency-relevant histone marks (e.g. H3K56ac) may be one mechanism through which pluripotency maintenance is affected in FACT-depleted cells (Ding et al., 2012; C. Jeronimo, Poitras, & Robert, 2019; Pardo et al., 2010; Y. Tan et al., 2013; Tettey et al., 2019; Xie et al., 2009). This shuffling of histone modifications likely disrupts recruitment of factors that maintain gene expression by sensing histone marks (e.g. recognition of methylated lysine residues on histones by CHD1 and CHD2). This disrupted factor recruitment and retention may explain many reductions in transcript abundance following FACT depletion. As FACT binding correlates with CHD1, CHD2, and gene expression and may remove CHD1 from partially unraveled nucleosomes (Farnung et al., 2021; Célia Jeronimo et al., 2020; Mylonas & Tessarz, 2018; Park, Shivram, & Iyer, 2014), CHD1 may also become trapped on chromatin without FACT-dependent displacement, thereby reducing expression of target genes.

RNAPII pausing is a phenomenon that occurs at the promoters of coding genes, as well as at eRNAs and PROMPTs (Gressel, Schwalb, & Cramer, 2019; Henriques et al., 2018; Tettey et al., 2019). As FACT has been shown to maintain pausing of RNAPII at coding promoters (Tettey et al., 2019), a plausible model emerges through which FACT represses transcription from these regions by maintaining RNAPII pausing to silence improper transcription. Given the enrichment of pluripotency- and differentiation-associated pathways found for the putative targets of these non-coding elements, this RNAPII pausing-mediated silencing may be the mechanism through which FACT prevents changes in cellular identity (i.e. reprogramming to iPSCs from fibroblasts) (Kolundzic et al., 2018; Mylonas & Tessarz, 2018; Shen et al., 2018; Tettey et al., 2019).

As many groups have suggested, the act of transcription by RNAPII itself may be responsible for destabilization of nucleosomes, creating a genomic conflict for FACT to resolve (Farnung et al., 2021; Formosa & Winston, 2020; Goswami et al., 2022; Célia Jeronimo et al., 2020; Y. Liu et al., 2020). With FACT depleted, this nucleosome destabilization likely compounds issues created by failure to maintain RNAPII pausing; it is likely that this combination of genome destabilization and failure to reassemble is responsible for the vast majority of derepressed transcription following FACT depletion. This model is further strengthened by a lack of FACT binding at putative silencers (Fig. S7A, C), and these regions do not display improper transcription after FACT depletion (Fig. S7B, D), suggesting that derepression by FACT depletion is not sufficient to induce transcription alone, but requires pre-initiated and paused RNAPII.

Together, the work presented here supports prior studies and enhances our understanding of the mechanistic role for FACT in mammalian pluripotent systems. Future work should aim to address the interplay between FACT, pluripotency factors, and histone modifications (such as H3K56ac), and the potential redistribution of modifications in contributing to alteration in cis-regulatory elements when FACT is lost or altered in disease settings.

## Author Contributions

D.C.K. and S.J.H. designed the study, wrote and edited the manuscript. D.C.K. performed most experiments. K.N.M. and S.J.H. generated cell lines. S.M.L. performed CUT&RUN experiments. D.C.K. analyzed the data with assistance from S.J.H.

## Acknowledgments

We thank members of the Hainer Lab for critical reading of the manuscript. We thank the ENCODE Consortium, the ENCODE production laboratories, and all other members of the scientific community who generated datasets that were essential to the completion of this study. This project used the NextSeq500 and NextSeq 2000 available at the University of Pittsburgh Health Sciences Sequencing Core at UPMC Children’s Hospital of Pittsburgh for sequencing with special thanks to its director, William MacDonald. This research was supported in part by the University of Pittsburgh Center for Research Computing through the resources provided. This work was supported by the Samuel and Emma Winters Foundation, 2018-2019 (to S.J.H.) and the National Institutes of Health Grant Number R35GM133732 (to S.J.H.).

## Materials and Methods

### Materials availability

Plasmids and cell lines generated in this study are available on request. All resources generated in this study must be acquired via a Material Transfer Agreement (MTA) granted by the University of Pittsburgh.

### Cell Lines

Mouse embryonic stem cells were derived from E14 (Hooper, Hardy, Handyside, Hunter, & Monk, 1987). Male E14 murine embryonic stem cells were grown in feeder-free conditions on 10 cm plates gelatinized with 0.2% porcine skin gelatin type A (Sigma) at 37°C and 5% CO_2_. Cells were cultured in Dulbecco’s Modified Eagle Medium (Gibco), supplemented with 10% Fetal Bovine Serum (Sigma, 18N103), 0.129mM 2-mercaptoethanol (Acros Organics), 2 mM glutamine (Gibco), 1X nonessential amino acids (Gibco), 1000U/mL Leukemia Inhibitory Factor (LIF), 3 µM CHIR99021 GSK inhibitor (p212121), and 1 µM PD0325091 MEK inhibitor (p212121). Cells were passaged every 48 hours using trypsin (Gibco) and split at a ratio of ∼1:8 with fresh medium. Routine anti-mycoplasma cleaning was conducted (LookOut DNA Erase spray, Sigma) and cell lines were screened by PCR to confirm no mycoplasma presence.

### Auxin Inducible Degradation

Cell lines were constructed in an E14 murine ES cell line with osTIR1 already integrated into the genome. SPT16 was C-terminally tagged using a 39 amino acid mini-AID construct also containing a 3xV5 epitope tag (Kubota, Nishimura, Kanemaki, & Donaldson, 2013; Natsume, Kiyomitsu, Saga, & Kanemaki, 2016; Nishimura, Fukagawa, Takisawa, Kakimoto, & Kanemaki, 2009; Nishimura & Kanemaki, 2014). Two homozygous isolated clones were generated using CRISPR-mediated homologous recombination with Hygromycin B drug selection and confirmed by PCR and Sanger sequencing.

Cells were depleted of AID-tagged SPT16 protein by addition of 500 nM 3-Indole Acetic Acid (3-IAA, Sigma) dissolved in 100% EtOH and pre-mixed in fresh medium. Cells were incubated with 3-IAA or 0.1% EtOH (vehicle) for 3, 6, 12, 16, or 24 hours to deplete the FACT complex and confirmed by Western blotting. Importantly, cells were cultured on 10 cm plates undisturbed for 48 hours prior to AID depletion, ensuring that relevant effects are not due to passaging-related disturbances.

### Alkaline Phosphatase Staining

Cells were treated with EtOH or 3-IAA as described above, with alkaline phosphatase staining after 6, 24, and 48 hours. Treated cells were washed twice in 1X Dulbecco’s Phosphate-Buffered Saline (DPBS, Gibco) and crosslinked in 1% formaldehyde (Fisher) in DPBS for five minutes at room temperature. Crosslinking was quenched with 500 mM glycine and cells were washed twice in 1xDPBS. Cells were stained with VECTOR Red Alkaline Phosphatase Staining Kit (Vector Labs) per manufacturer’s instructions in a 200 mM Tris-Cl buffer, pH 8.4. 8 mL working solution was added to each 10 cm plate and incubated in the dark for 30 minutes before being washed with DPBS and imaged.

### Western blotting

Western blotting was performed using a mouse monoclonal anti-V5 epitope antibody (Invitrogen 46-0705, lot 1923773), a mouse monoclonal anti-SSRP1 antibody (BioLegend 609702, lot B280320), and a mouse monoclonal anti-beta-actin loading control (Sigma). Secondary antibody incubations were performed with goat polyclonal antibodies against either rabbit or mouse IgG, (BioRad 170-6515, lot #64149722, BioRad 170-6516, lot #64147779). Crude protein extractions were performed using RIPA buffer (150 mM NaCl, 1% IPEGAL CA-630, 0.5% sodium deoxycholate, 0.1% sodium dodecyl sulfate, 25 mM Tris-Cl, pH 7.4) with freshly added protease inhibitors (ThermoFisher) and flash-frozen immediately after extraction. Samples were quantitated using the Pierce BCA Protein Assay kit (ThermoFisher). 20 µg were diluted in RIPA buffer with 10 mM dithiothreitol (DTT) and Laemmeli sample buffer before being loaded on 7.5% Tris-acrylamide gels for Western blotting. Proteins were transferred to nitrocellulose membranes (BioTrace) via a Criterion tank blotter (BioRad) at 100V for one hour and stained with 0.5% Ponceau S (Sigma) in 1% acetic acid to confirm proper transfer. Membranes were blocked in 5% milk in PBST prior to overnight primary antibody incubation at 4°C. Membranes were then washed and incubated in secondary antibody (Bio-Rad) for one hour at room temperature, washed, and developed with SuperSignal West Pico chemiluminescent reagent (ThermoFisher) for 5 minutes at room temperature.

### CUT&RUN

CUT&RUN was performed as described (Hainer et al., 2019; Hainer & Fazzio, 2019; Patty & Hainer, 2021; Skene & Henikoff, 2017), using recombinant Protein A/Protein G-MNase (pA/G-MN) (Meers, Bryson, et al., 2019). Briefly, 100,000 nuclei were isolated from cell populations using a hypotonic buffer (20 mM HEPES-KOH, pH 7.9, 10 mM KCl, 0.5mM spermidine, 0.1% Triton X-100, 20% glycerol, freshly added protease inhibitors) and bound to lectin-coated concanavalin A magnetic beads (200 µL bead slurry per 500,000 nuclei) (Polysciences). Immobilized nuclei were chelated with blocking buffer (20 mM HEPES, pH 7.5, 150 mM NaCl, 0.5mM spermidine, 0.1% BSA, 2mM EDTA, fresh protease inhibitors) and washed in wash buffer (20 mM HEPES, pH 7.5, 150 mM NaCl, 0.5mM spermidine, 0.1% BSA, fresh protease inhibitors). Nuclei were incubated in wash buffer containing primary antibody (anti-V5 mouse monoclonal, Invitrogen 46-0705, lot 1923773) for one hour at room temperature with rotation, followed by incubation in wash buffer containing recombinant pA/G-MN for 30 minutes at room temperature with rotation. Controls lacking a primary antibody were subjected to the same conditions but incubated in wash buffer without antibody prior to incubation with pA/G-MN. Samples were equilibrated to 0°C and 3 mM CaCl_2_ was added to activate pA/G-MN cleavage. After suboptimal digestion for 15 minutes, digestion was chelated with 20 mM EDTA and 4 mM EGTA, and 1.5 pg MNase-digested *S. cerevisiae* mononucleosomes were added as a spike-in control. Genomic fragments were released after an RNase A treatment. After separating released fragments through centrifugation, fragments isolated were used as input for a library build consisting of end repair and adenylation, NEBNext stem-loop adapter ligation, and subsequent purification with AMPure XP beads (Agencourt). Barcoded fragments were then amplified by 14 cycles of high-fidelity PCR and purified using AMPure XP. Libraries were pooled and sequenced on an Illumina NextSeq500 to a depth of ∼10 million mapped reads.

### CUT&RUN data analysis

Paired-end fastq files were trimmed to 25 bp and mapped to the mm10 genome with bowtie2 (options -q -N 1 -X 1000) (Langmead & Salzberg, 2012). Mapped reads were duplicate-filtered using Picard (“Picard Tools, Broad Institute,”) and filtered for mapping quality (MAPQ ≥ 10) using SAMtools (H. Li et al., 2009). Size classes corresponding to FACT footprints (<120 bp) were generated using SAMTools (H. Li et al., 2009). Reads were converted to bigWig files using deepTools (options -bs 1 --normalizeUsing RPGC, --effectiveGenomeSize 2862010578) (Ramirez, Dundar, Diehl, Gruning, & Manke, 2014), with common sequencing read contaminants filtered out according to ENCODE blacklisted sites for mm10. Heatmaps were generated using deepTools computeMatrix (options -a 2000 -b 2000 -bs 20 --missingDataAsZero) and plotHeatmap (Ramirez et al., 2014). Peaks were called from CUT&RUN data using SEACR, a CUT&RUN-specific peak-calling algorithm with relaxed stringency and controls lacking primary antibody used in lieu of input data (Meers, Bryson, et al., 2019). Motifs were then called from these peaks using HOMER with default settings (Heinz et al., 2010). Pathway analysis was performed on peaks present in at least 2/4 SPT16-V5 CUT&RUN experiments using HOMER and the WikiPathways database, then plotted in GraphPad Prism 10, with the y-axis representing rank of enrichment (Heinz et al., 2010).

One-dimensional heatmaps were generated by the same pipeline for CUT&RUN and ChIP-seq data. Matrices generated using deepTools computeMatrix as above were averaged by position relative to reference point using plotProfile with the option –outFileNameMatrix. Average position scores per technical replicate were then averaged together and translated to colorimetric scores using ggplot2.

### Transient Transcriptome Sequencing

TT-seq was performed using a modified method (Dolken et al., 2008; Duffy et al., 2015; Radle et al., 2013; Schwalb et al., 2016). 500 mM 4sU (Carbosynth T4509) was dissolved in 100% DMSO (Fisher). Following protein depletion as above, cells were washed with 1x DPBS (Corning), resuspended in medium containing 500 µM 4sU, and incubated at 37°C and 5% CO_2_ for five minutes to label nascent transcripts. After washing cells with 1x DPBS, RNA was extracted with TRIzol and fragmented using a Bioruptor Pico for one cycle at high power. Thiol-specific biotinylation of 100 g of total RNA was carried out using 10x biotinylation buffer (100 mM Tris-Cl, pH 7.4, 10 mM ethylenediaminetetraacetic acid) and EZ-Link Biotin-HPDP (Pierce 21341) dissolved in dimethylformamide at 1 mg/mL. Biotinylation was carried out for 2 hours away from light with 1000 rpm shaking at 37°C. RNA was extracted with chloroform and precipitated using NaCl and isopropanol. Labeled RNA was separated from unlabeled RNA via a streptavidin C1 bead-based pulldown (DynaBeads, ThermoFisher). In brief, beads were washed in bulk in 1 mL of 0.1N NaOH with 50mM NaCl, resuspended in binding buffer (10mM Tris-Cl, pH 7.4, 0.3M NaCl, 1% Triton X-100) and bound to RNA for 20 minutes at room temperature with rotation. Beads bound to labeled RNA were washed twice with high salt wash buffer (5 mM Tris-Cl, pH 7.4, 2M NaCl, 1% Triton X-100), twice with binding buffer, and once in low salt wash buffer (5 mM Tris-Cl, pH 7.4., 1% Triton X-100). Nascent RNA was recovered from beads using two elutions with fresh 100mM dithiothreitol at 65°C for five minutes with 1000 rpm shaking. Recovered nascent RNA was then extracted with PCI and chloroform, and then isopropanol precipitated.

Strand-specific nascent RNA-seq libraries were built using the NEBNext Ultra II Directional Library kit, with the following modifications: 200 ng of fragmented RNA was used as input for ribosomal RNA removal via antisense tiling oligonucleotides and digestion with thermostable Rnase H (MCLabs) (Adiconis et al., 2013; Morlan, Qu, & Sinicropi, 2012). rRNA-depleted RNA samples were treated with Turbo DNase (ThermoFisher) and purified by silica column (Zymo RNA Clean & Concentrator). RNA was fragmented at 94°C for five minutes and subsequently used as input for cDNA synthesis and strand-specific library building according to manufacturer protocol. Libraries were pooled and sequenced via Illumina NextSeq500 or NextSeq2000 to a sequencing depth of a minimum of 40 million mapped reads.

### TT-seq data analysis

Paired-end fastq files were trimmed and filtered using Trim Galore (Krueger, 2015), then aligned to the mm10 mouse genome using STAR (options --outSAMtype SAM -- outFilterMismatchNoverReadLmax 0.02 --outFilterMultimapNmax 1). Feature counts were generated using subread featureCounts (options -s 2 -p -B) for genes, PROMPTs, DHSs, and putative eRNAs based on genomic coordinates (see next paragraph). Reads were imported to R and downstream analysis was conducted using DESeq2 (Love, Huber, & Anders, 2014). Differentially expressed transcripts were plotted using EnhancedVolcano (Blighe K, 2021). Pathway analysis was performed on all significantly up- and downregulated genes separately using HOMER with the WikiPathways database (Heinz et al., 2010). Significance was defined as DESeq2 adjusted p-value < 0.05. Top five enriched categories were plotted in GraphPad Prism 10 against -log_10_ p-value, with manually curated categories added from the top 50 hits. Y-axes indicate pathway enrichment ranking. For downstream analyses, we generated GTF and bed files of Gencode mm10 vM25 genes, sorted by nascent transcription in all control (Untagged, 0h, and EtOH-treated) samples, pooled together.

Non-coding transcripts were identified by removing all transcription start sites within 1kb of annotated mm10 coding genes from the previously described gene-distal DNaseI hypersensitive sites (GSM1014154) (Consortium, 2012; Davis et al., 2018; Thurman et al., 2012). PROMPTs were called by genomic location (within 1 kb of an annotated mm10 TSS and divergently transcribed to the TSS). ncRNAs were assigned to the closest coding gene and pathway analysis was conducted as above. Putative enhancers were defined as overlapping a DHS, as well as the presence of either H3K27ac or H3K4me1, according to ChIP-seq data from ENCODE (Consortium, 2012; Davis et al., 2018; Thurman et al., 2012).

### Reverse Transcription and quantitative PCR (RT-qPCR)

RT-qPCR was performed as previously described (Hainer et al., 2015). Briefly, RNA was extracted from cells using TRIzol following treatment with either 3-IAA or EtOH for 0, 3, and 6 hours. 1 µg of RNA was used as input for reverse transcription, and quantitative PCR was performed using 5 µM PCR primers targeting the gene of interest with KAPA SYBR green master mix. Technical replicates shown represent the average of three individual qPCR reactions for each treatment/target/condition group. Error bars shown represent the standard deviation of two replicates for each combination.

### Assay for Transposase-Accessible Chromatin Sequencing (ATAC-seq)

Omni-ATAC-seq was performed as previously described (Corces et al., 2017). Briefly, cells were depleted of FACT proteins using a treatment with EtOH (vehicle) or 500 µM 3-IAA for 0, 3, 6, 12, or 24 hours. Nuclei were extracted from 60,000 cells as described for CUT&RUN and flash-frozen until use. Frozen nuclei were resuspended in transposition mix containing 1X TD buffer (10 mM Tris pH 7.6, 5 mM MgCl_2_, 10% dimethylformamide), DPBS, 0.1% Tween-20, 1% digitonin, and 4 µL Tn5 transposome (Diagenode) per reaction. Samples were incubated at 37°C for 30 minutes with 1000 rpm shaking. Transposed DNA was purified using a Clean and Concentrator kit (Zymo) per manufacturer’s instructions. Samples were amplified for 5 cycles of high-fidelity PCR (KAPA), then held on ice and assessed via qPCR (KAPA SYBR Green). Samples were then returned to the thermocycler for as many cycles as needed to reach 1/3 qPCR saturation (∼10 total cycles). Amplified libraries were gel-extracted between 150-500 bp and sequenced via Illumina NextSeq2000 to a sequencing depth of ∼50 million mapped reads.

### ATAC-seq data analysis

Paired-end fastq files were trimmed to 25 bp and mapped to the mm10 genome with Bowtie 2 (using the options --very-sensitive --dovetail -q -N 1 -X 1000) (Langmead & Salzberg, 2012). Mapped reads were duplicate filtered using Picard (“Picard Tools, Broad Institute,”) and filtered for mapping quality (MAPQ ≥ 10) using SAMtools (H. Li et al., 2009). Reads were separated into size classes of 1-100 bp (factor binding) and 180-247 bp (mononucleosomal fragments) using an awk command. Size-selected reads were converted to bigWig files using deepTools (options -bs 1 --normalizeUsing RPGC, --effectiveGenomeSize 2308125349 -- ignoreForNormalization chrM -e) (Ramirez et al., 2014). Differential bigwigs were generated using deepTools bigwigCompare (-bs 10) (Ramirez et al., 2014). Heatmaps were generated using deepTools computeMatrix (options --referencePoint TSS -a 2000 -b 2000 -bs 20 -- missingDataAsZero) and plotHeatmap, based on the 1-100 size class (Ramirez et al., 2014). Differences in accessibility were plotted by generating matrices in deepTools as above. Where indicated, data were clustered using k-means clustering.

### Micrococcal Nuclease Sequencing (MNase-seq)

MNase-seq was performed as previously described (Hainer et al., 2015). In brief, cells were depleted of FACT proteins using a 24-hour treatment with EtOH (vehicle) or 500 nM 3-IAA, 5 million cells were collected, crosslinked using 1% formaldehyde for 15 minutes at RT, and quenched with 500 mM glycine. Cells were lysed in hypotonic buffer (10 mM Tris-Cl, pH 7.5, 10 mM NaCl, 2 mM MgCl_2_, 0.5% NP-40, 0.3 mM CaCl_2_, and 1X protease inhibitors) and subjected to 5 minutes of digestion with MNase (TaKaRa) at 37°C before chelation with EDTA and EGTA. Samples were treated with Rnase A (ThermoFisher) for 40 minutes at 37C and 1000 rpm constant shaking in a thermomixer. Crosslinks were reversed overnight at 55°C and chromatin was digested with Proteinase K, then used as input for a paired-end library build.

1 µg input DNA was treated with Quick CIP (NEB) for 30 minutes and heat-inactivated. End repair was then performed using T4 DNA Polymerase (NEB), T4 Polynucleotide Kinase (NEB), and Klenow DNA Polymerase (NEB) simultaneously. A-overhangs were added to sequences via treatment with Klenow Polymerase without exonuclease activity and Illumina paired-end TruSeq adapters were added using Quick Ligase (NEB). Barcoded DNA was purified using AMPure XP beads (Agencourt) and amplified by high-fidelity PCR (KAPA). Completed libraries were subjected to silica column purification (Zymo DNA Clean & Concentrator) and sequenced via Illumina NextSeq500 to a sequencing depth of ∼50 million mapped reads.

### MNase-seq data analysis

Paired-end fastq files were trimmed to 25 bp and mapped to the mm10 genome with bowtie2 (using the options -q -N 1 -X 1000) (Langmead & Salzberg, 2012). Mapped reads were duplicate-filtered using Picard (“Picard Tools, Broad Institute,”) and filtered for mapping quality (MAPQ ≥ 10) using SAMtools (H. Li et al., 2009). Reads were then sorted into nucleosome- (135-165 bp), subnucleosome- (100-130 bp), and transcription factor- (<80 bp) sized fragments using SAMtools (H. Li et al., 2009). Nucleosome-sized reads were converted to bigWig files using deepTools (options -bs 1 --normalizeUsing RPGC, --effectiveGenomeSize 2862010578), with common sequencing read contaminants filtered out according to ENCODE blacklisted sites for mm10 (Ramirez et al., 2014). Differential bigwigs were generated using deepTools bigwigCompare (default options) (Ramirez et al., 2014). Heatmaps were generated using deepTools computeMatrix (options --referencePoint TSS -a 2000 -b 2000 -bs 20 -- missingDataAsZero) and plotHeatmap and plotProfile (Ramirez et al., 2014). Differences in nucleosome occupancy were plotted by generating matrices in deepTools as above. Metaplots of MNase-seq data in Figures 7 and S8 include standard error shaded around the plotted line (mean).

### Statistics

Statistical details for each experiment shown can be found in the accompanying figure legends. Where indicated, “n” designates independent technical replicates for the same biological sample, while biological replicates are referred to as “clone 1” and “clone 2” to differentiate between independently targeted cell lines. Statistical tests were used in TT-seq analyses as per the default parameters for DESeq2, with a correction applied to minimize fold change of lowly-expressed transcripts (LFCshrink), as well as motif analysis (default HOMER parameters) and peak-calling (default SEACR and HOMER parameters for CUT&RUN and ChIP-seq datasets, respectively). Any error bars shown represent one standard deviation in both directions. Standard error was calculated via deepTools plotProfile for MNase-seq metaplots generated in Figures 7 and S8. Significance was defined as a p-value < 0.05 by the respective test performed (indicated with “*”). No data or subjects were excluded from this study. Average values for CUT&RUN, ChIP-seq, and MNase-seq datasets were determined by computing the mean of coverage at each base pair throughout the genome between replicates. Merged replicates indicates mean of read-coverage normalized tracks generated for each individual replicate.

## Data availability

This paper analyzes existing, publicly available data housed in the NCBI Gene Expression Omnibus (GEO) and the Sequence Read Archive (SRA). The accession numbers for the datasets are listed throughout the manuscript. Unedited raw sequencing reads and processed bigwig files generated during this study have been deposited in NCBI GEO and the SRA and will be made public at time of formal publication. Any additional information required regarding the data reported in this paper is available from the lead contact upon request.

## Supplementary Information

**Supplementary Table 1.**
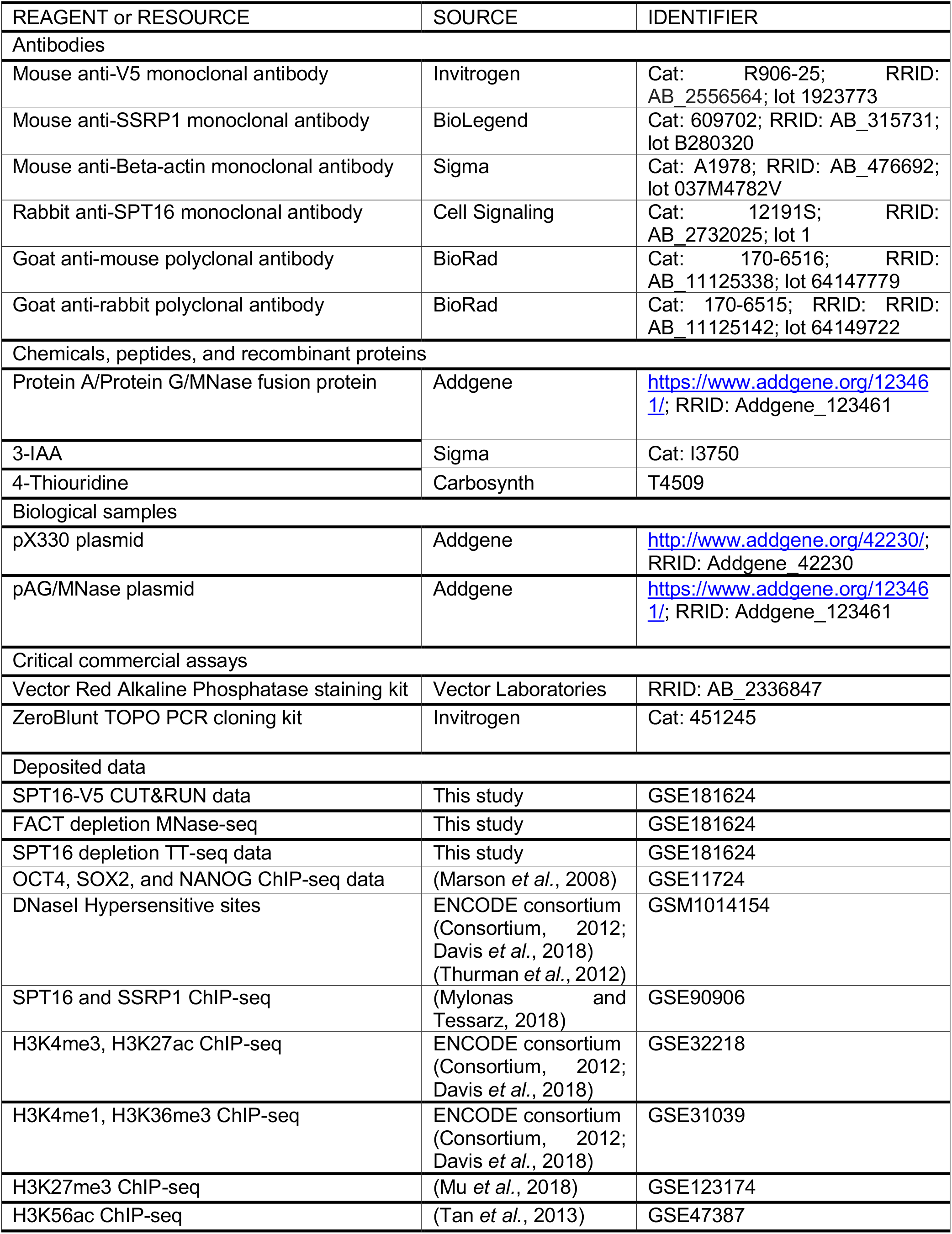
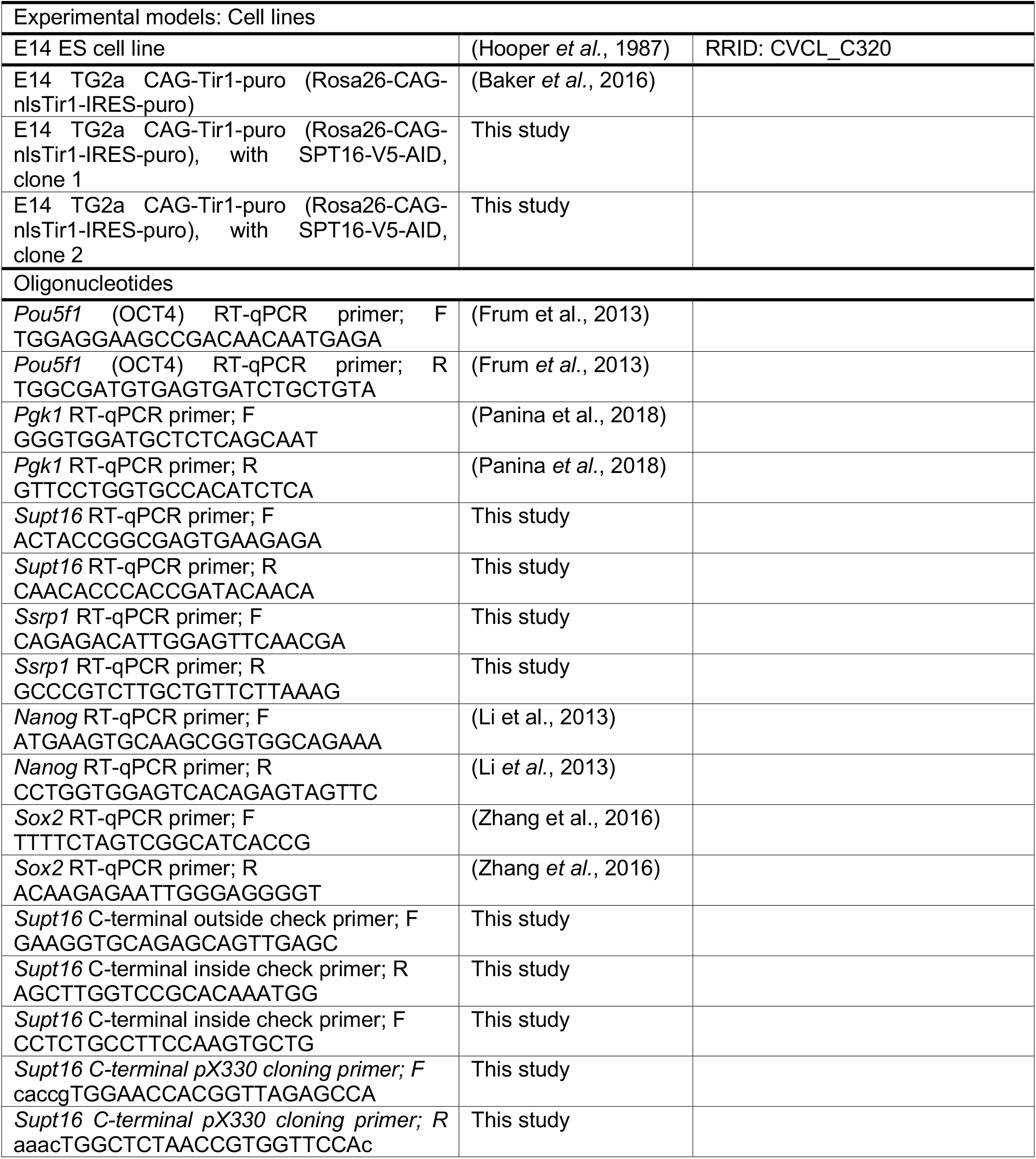
Key reagents, cell lines, and datasets used in this work.

## Supplemental Figures

**Fig. S1.**
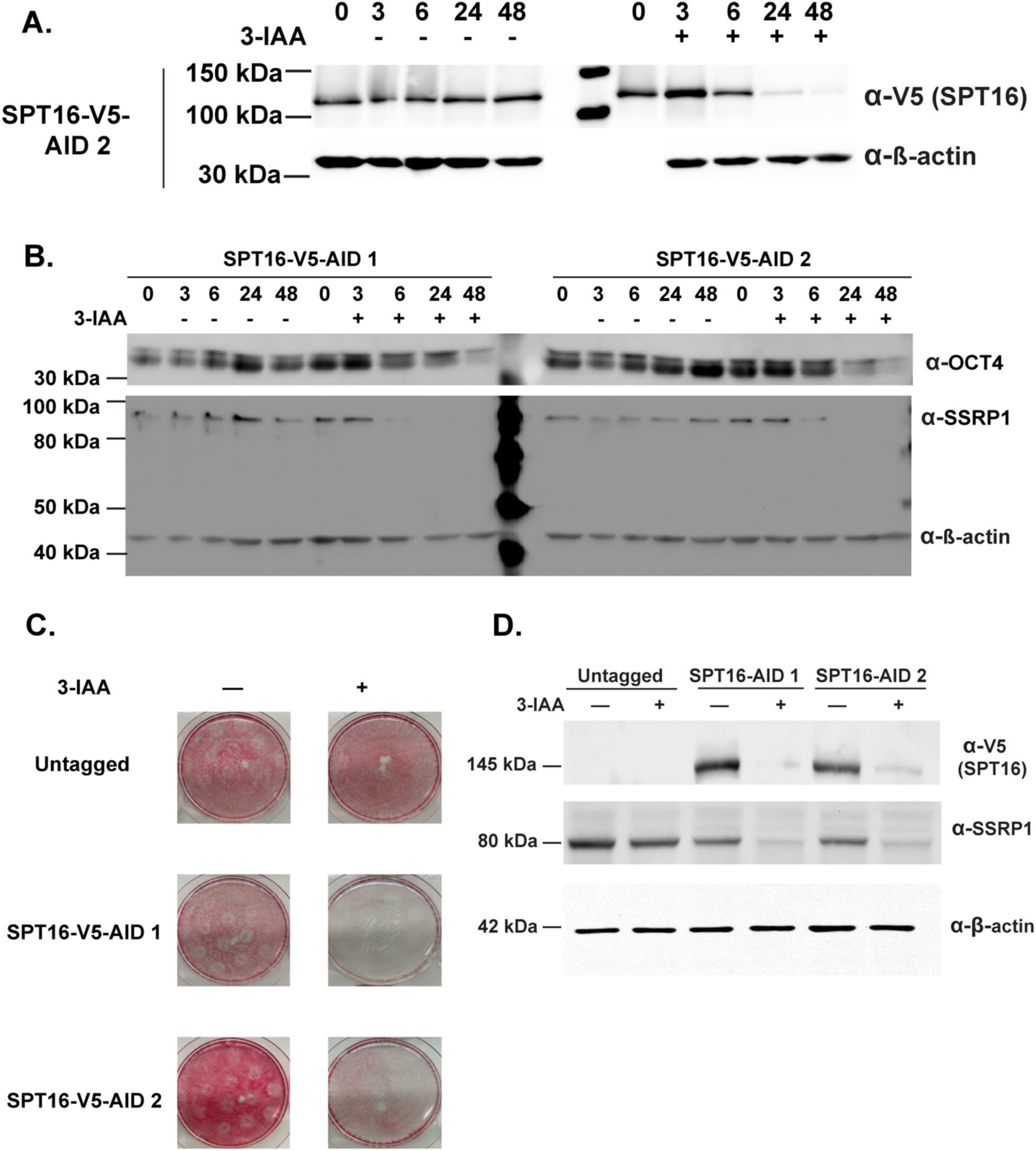
(Related to Fig. 1). Characterization of SPT16-V5-AID cell lines. A. Timecourse of 3-IAA treatment for SPT16 depletion in two independently targeted SPT16-V5-AID cell lines. 40 µg total protein loaded. Top to bottom, anti-V5 (targeting SPT16), anti-β-actin, anti-V5 (targeting SPT16), and anti-β-actin. 3-IAA – indicates vehicle control (EtOH). B. Timecourse of OCT4 (top), SSRP1 (middle) and β-actin (bottom) protein levels following 3-IAA treatment to deplete SPT16. 40 µg total protein loaded. 3-IAA – indicates vehicle control (EtOH). C. 10 cm plate images of alkaline phosphatase-stained cells following 24 hours of 3-IAA treatment (right) or vehicle control (left).

**Fig. S2.**
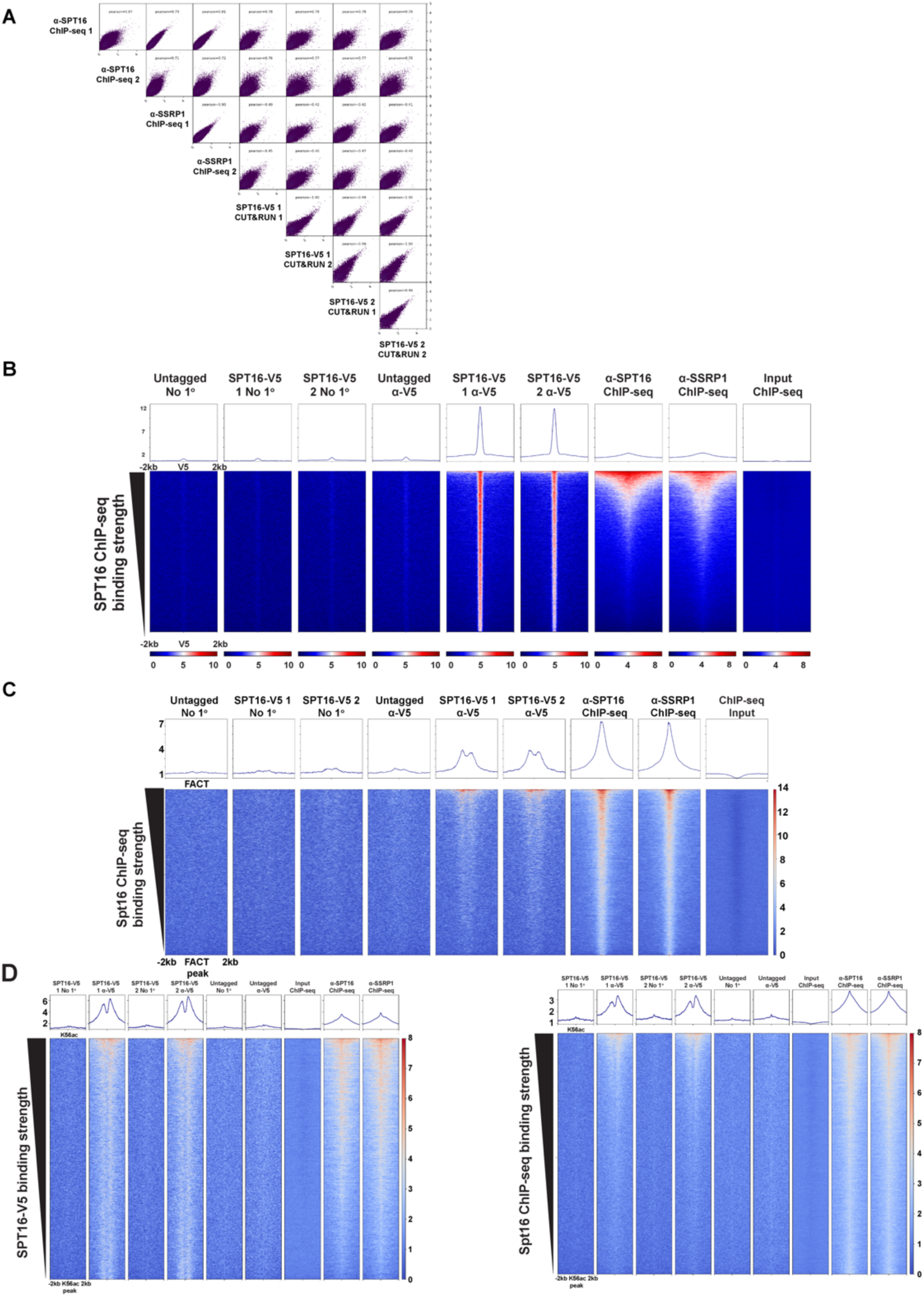
(Related to Fig. 2). Validation of SPT16-V5 CUT&RUN data. A. Pairwise scatterplots showing Pearson correlation between SPT16-V5 CUT&RUN, SPT16 ChIP-seq, and SSRP1 ChIP-seq. Individual technical replicates are compared for each sample. Bins represent average coverage over 5kb regions of the genome. B. SPT16-V5 CUT&RUN and published SPT16 and SSRP1 ChIP-seq data visualized over peaks called from SPT16-V5 CUT&RUN data using SEACR (ChIP-seq data: GSE90906) (Mylonas and Tessarz, 2018). C. SPT16-V5 CUT&RUN and published SPT16 and SSRP1 ChIP-seq data visualized over peaks called from SPT16 and SSRP1 ChIP-seq data using HOMER (ChIP-seq data: GSE90906) (Mylonas and Tessarz, 2018). D. FACT profiling data visualized at H3K56ac ChIP-seq peaks, +/− 2kb. Left heatmaps are visualized at SPT16-V5 CUT&RUN-bound peaks called from H3K56ac ChIP-seq data, while right heatmaps are visualized at FACT ChIP-seq bound peaks called from H3K56ac ChIP-seq data.

**Fig. S3.**
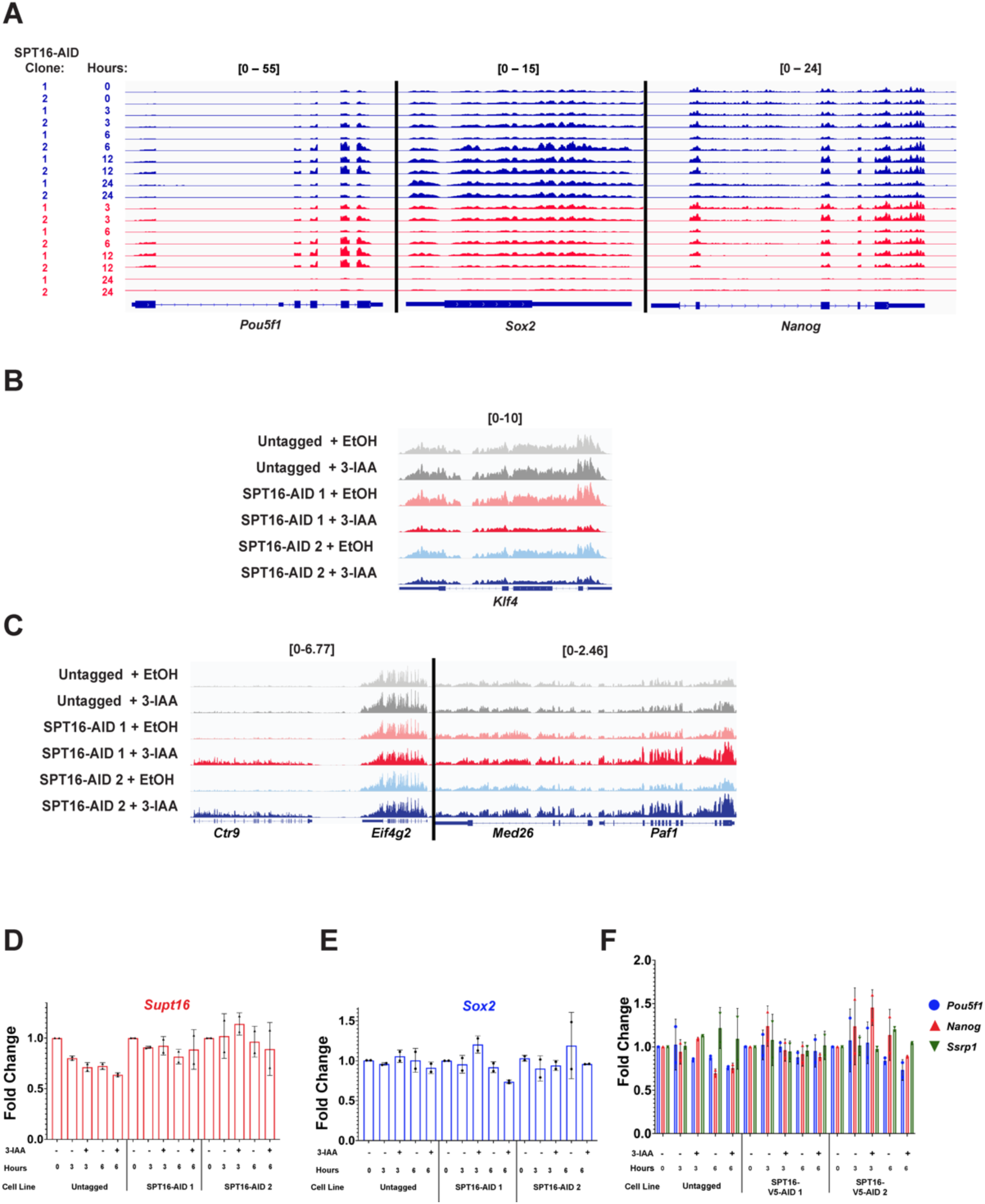
(Related to Fig. 3). Characterization of transcriptomic effects of SPT16 depletion and FACT interactions with pluripotency factors. A. IGV genome browser tracks depicting nascent transcription at the *Pou5f1* (left), *Sox2* (middle), and *Nanog* (right) genomic loci. Samples were treated with either 3-IAA (red) or vehicle (blue) for the indicated length. 24-hour samples are averaged technical replicates (n = 3); all other samples are individual technical replicates. B. IGV genome browser tracks depicting nascent transcription at the *Klf4* (right) genomic locus following 24 hours of 3-IAA treatment to deplete SPT16. Browser tracks represent merged technical replicates (n = 3), while biological replicates are displayed separately. C. As in B but depicting nascent transcription at the *Ctr9* (left) and *Paf1* (right) genomic loci. Technical replicates are averaged (n = 3). D-F. Short-term 3-IAA treatment (3- and 6-hour) for SPT16 depletion followed by RT-qPCR. Fold change calculated using ΔΔCt with normalization to *Pgk1* transcript abundance, where 0h timepoint is set to 1 and other timepoints are made relative. Error bars represent one standard deviation of fold change (n = 2 biological replicates). D. *Spt16* mRNA abundance. E. *Sox2* mRNA abundance. F. *Pou5f1* (blue), *Nanog* (red), and *Ssrp1* (green) mRNA abundances.

**Fig. S4.**
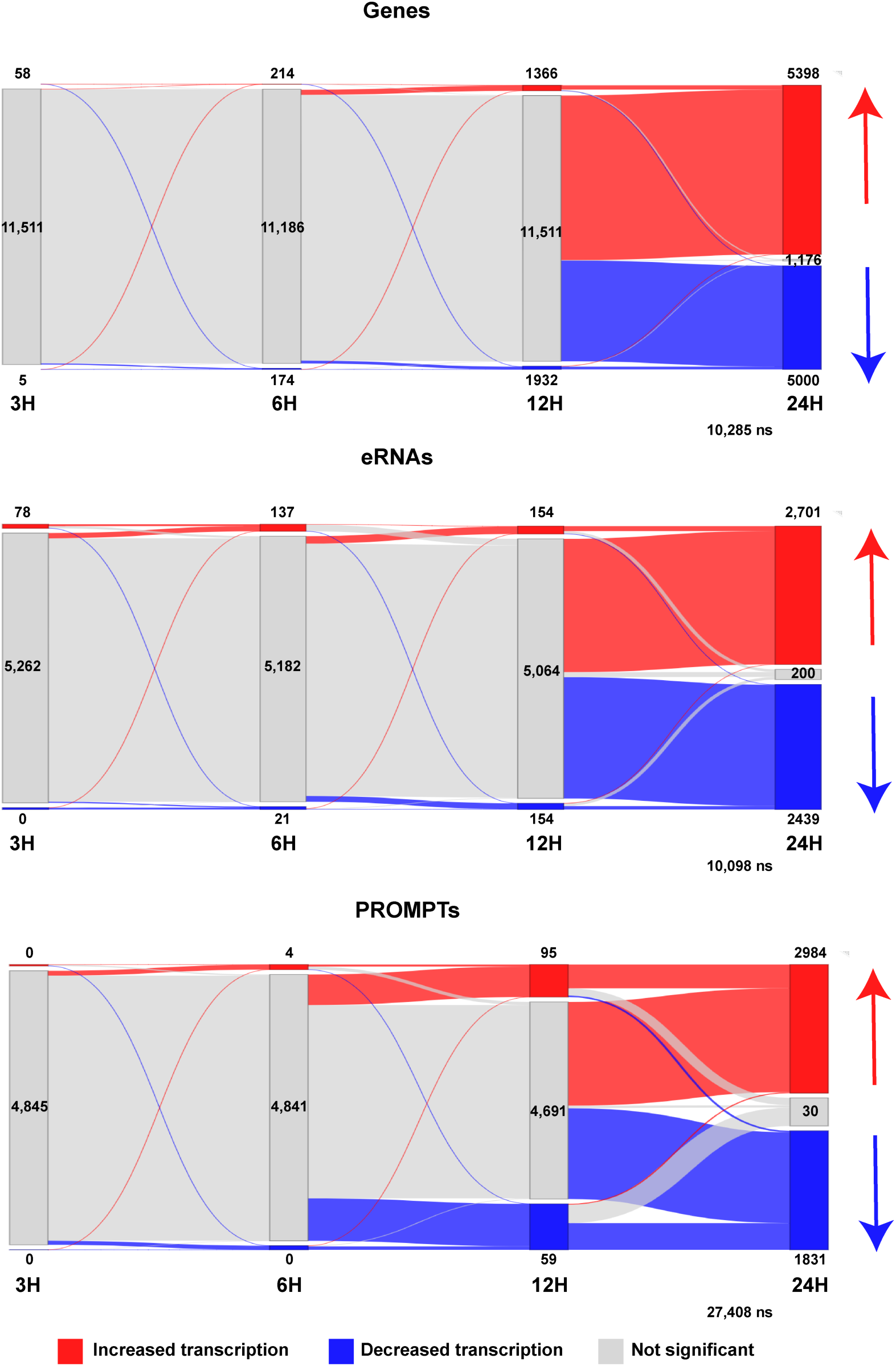
(Related to Figs. 3 and 4). Sankey plots depicting altered transcripts at 3, 6, 12, and 24 hours of treatment. Transcripts which never significantly changed were not plotted (ns). Red indicates increased transcripts, while blue indicates decreased transcriptions between timepoints. Each node indicates transcripts in one category at one timepoint, while flows indicate transcript changes between timepoints. Input values were taken from DESeq2 results listed in Table 1.

**Fig. S5.**
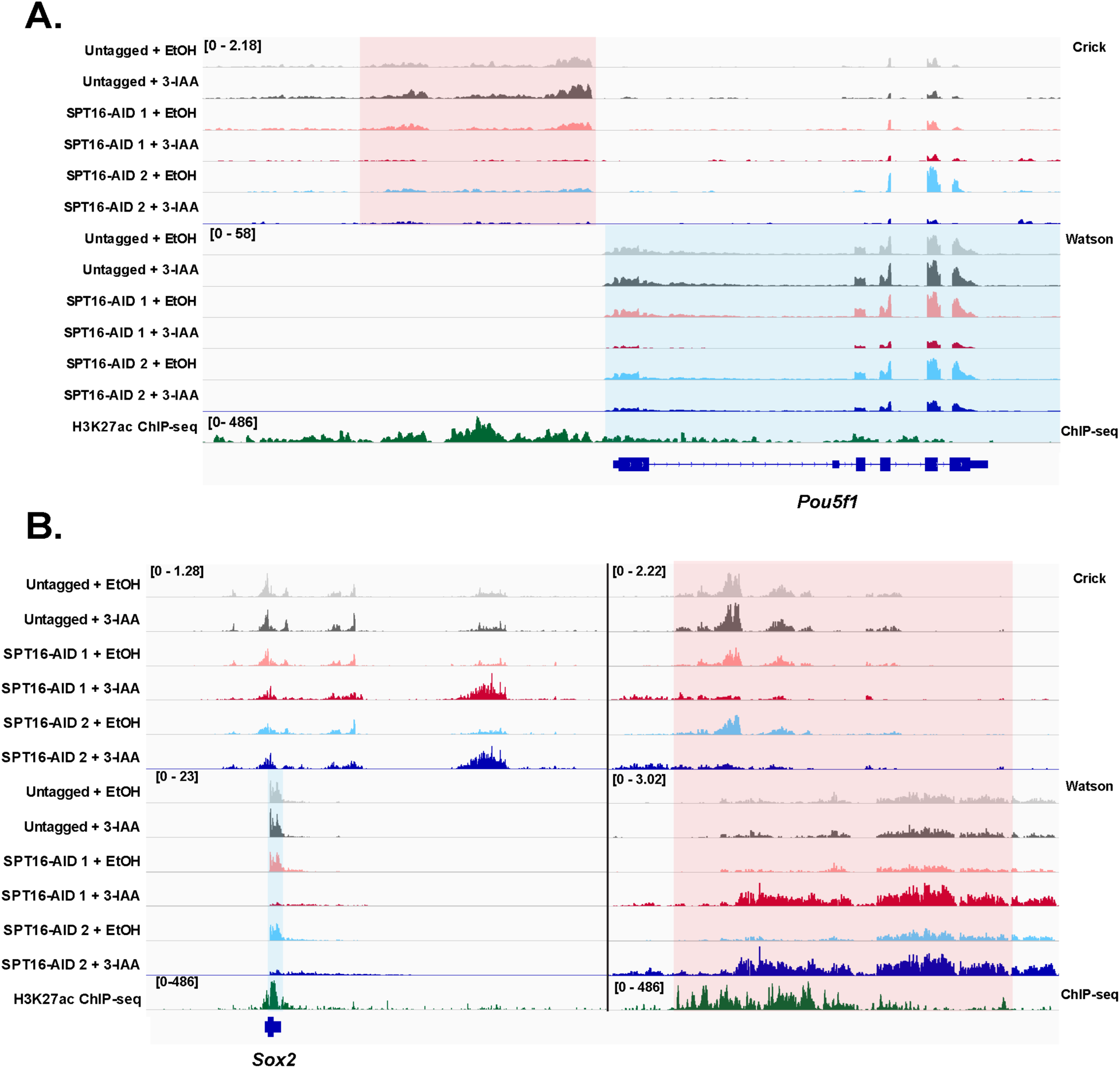
(Related to Fig. 4). FACT depletion reduces transcription at superenhancers. A. IGV genome browser tracks depicting nascent transcription at the *Pou5f1* locus, along with published H3K27ac ChIP-seq data. Red shaded area denotes a proximal superenhancer of *Pou5f1* transcription, while blue shaded area denotes the *Pou5f1* gene. Browser tracks represent merged technical replicates (n = 3), while biological replicates are displayed separately. B. IGV genome browser tracks depicting nascent transcription at the *Sox2* locus. Browser tracks represent merged technical replicates (n = 3), while biological replicates are displayed separately. Two individually scaled windows are shown to highlight eRNA transcription from the *Sox2* distal superenhancer (red shaded area) and nascent transcription from the *Sox2* genomic locus (blue shaded area).

**Fig. S6.**
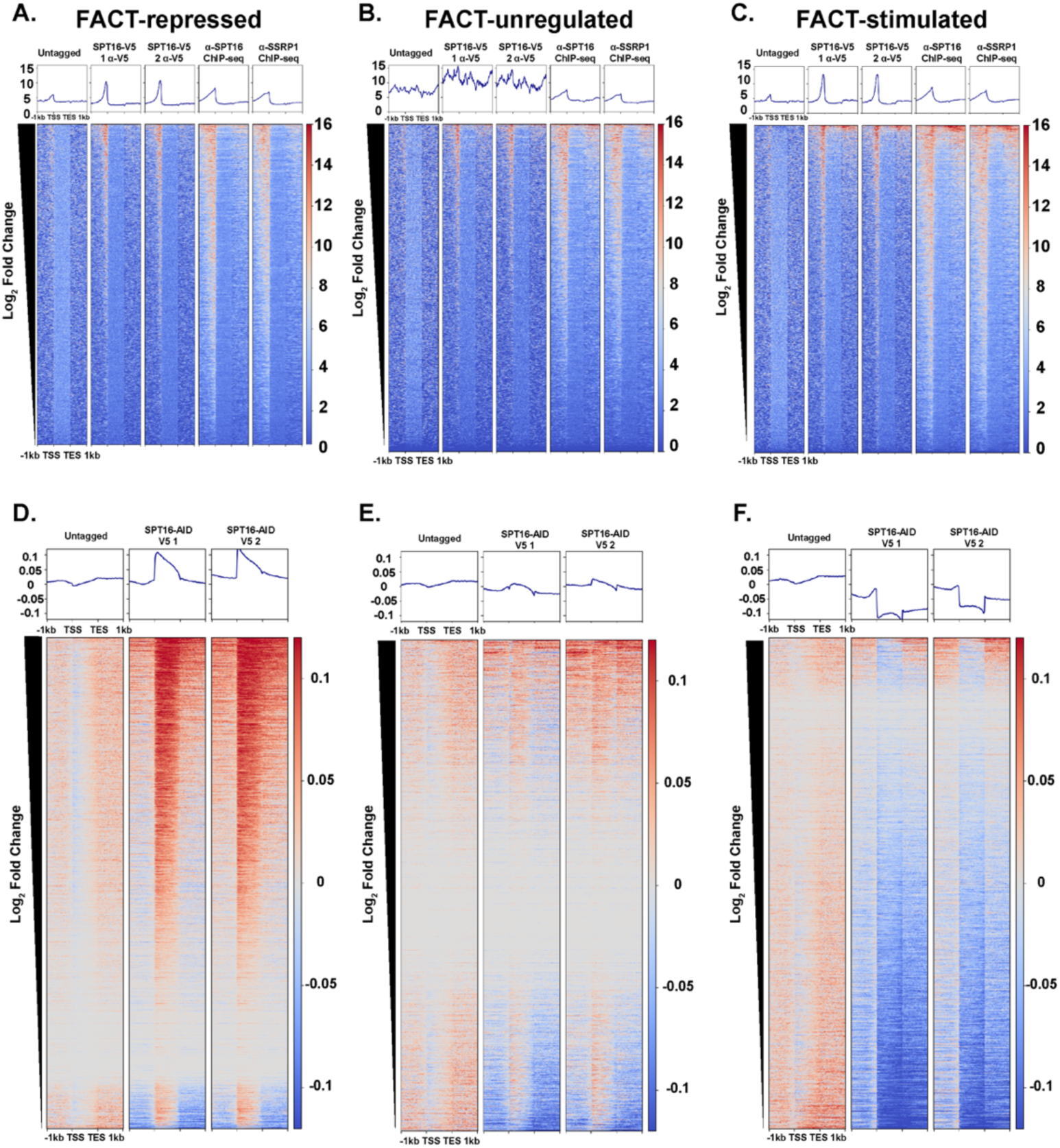
(Related to Fig. 4). SPT16-V5 binding is enriched at promoters of FACT-regulated genes. A-C. SPT16-V5 CUT&RUN and FACT ChIP-seq, visualized over genes classified by transcriptional change after 24 hours of 3-IAA treatment to deplete SPT16. Merged replicates shown as metagene plots, +/− 1kb from the start or end site of transcription (N = 3 for untagged, n = 2 for all other samples). Genes sorted by descending log2 fold change in DESeq2 results. Visualized over genes with significantly increased transcription (padj < 0.05 log2 fold change > 0.75) (A), genes with expression unaffected after FACT depletion (padj > 0.05 or log2 fold change < 0.75) (B), and genes with reduced transcription following FACT depletion (padj < 0.05, log2 fold change > 0.75) (C). D-F. Nascent transcription following 24 hours of 3-IAA treatment to deplete SPT16. Merged replicates shown as metagene plots, +/− 1 kb from the start or end site of transcription (n = 3). Genes sorted by descending log2 fold change in DESeq2 results. Visualized over significantly increased transcription (D), unchanged transcription (E), or reduced transcription (F) as in A-C.

**Fig. S7.**
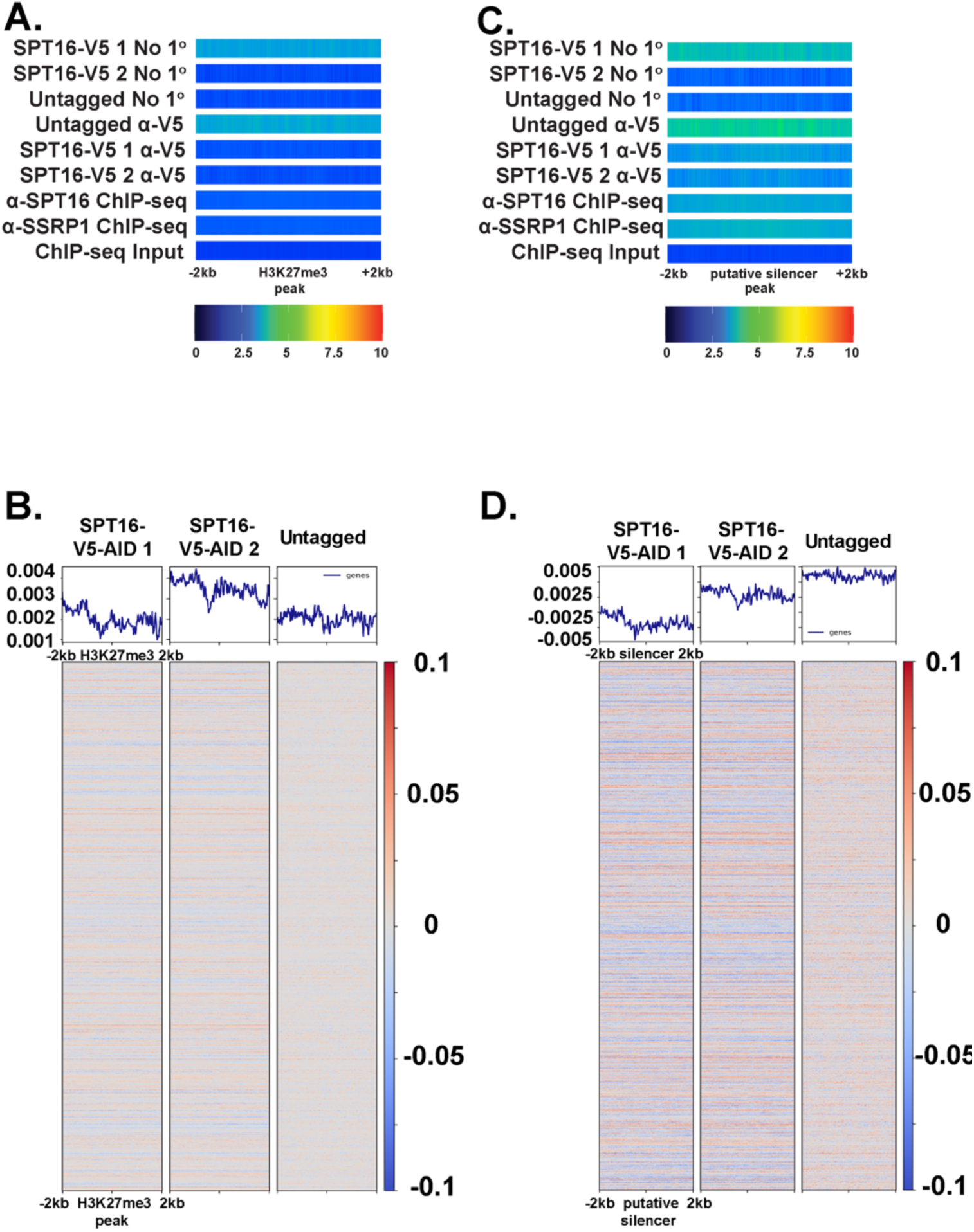
(Related to Figs 6 and 7) FACT neither binds at nor regulates transcription from regions marked by H3K27me3. A. FACT CUT&RUN (SPT16-V5) and ChIP-seq (GSE90906) data visualized at H3K27me3 ChIP-seq peaks +/− 2kb as one-dimensional heatmaps (K27me3 ChIP-seq from GSE123174; as in Fig. 5) (Mu et al., 2018; Mylonas and Tessarz, 2018). Shown as average of technical replicates, while biological replicates are displayed separately (n = 3 for for untagged CUT&RUN, n = 2 for all other samples). B. TT-seq data visualized at H3K27me3 ChIP-seq peaks +/− 2kb. Merged replicates shown as ratio of transcription in 3-IAA-treated samples to EtOH-treated samples (n = 3). C. As in S7A but visualized over putative silencers (defined as gene-distal DHSs overlapping an H3K27me3 peak). D. As in S7B but visualized over putative silencers.

**Fig. S8.**
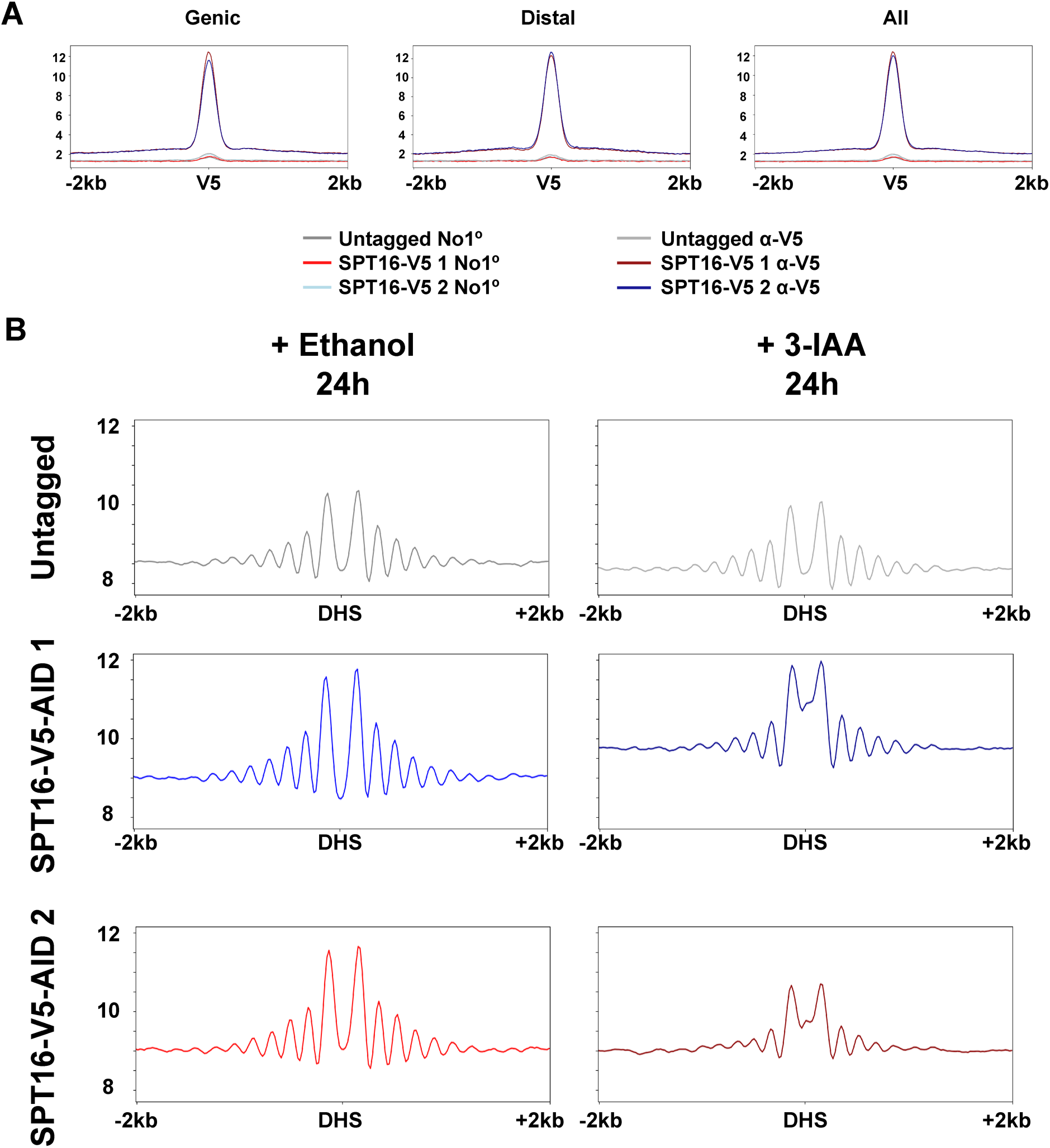
(Related to Figs. 6-7). FACT regulates genic and gene-distal binding sites similarly. A. Metaplots depicting SPT16-V5 binding over SPT16-V5 binding sites. Average signal over V5 sites shown with standard error shaded. Sites overlapping promoters (left), not overlapping genes (middle) and all together (CUT&RUN data averaged as in Fig. 2; n = 3 for untagged samples, n = 2 for others). B. Metaplots of MNase-seq data following 24 hours of SPT16 depletion, visualized over gene-distal DHSs. Metaplots shown represent merged technical replicates, while biological replicates are shown separately (n = 3 for untagged, n = 2 for each AID-tagged clone). MNase-seq data visualized over gene-distal DNaseI hypersensitive sites, +/− 2kb (DNase-seq from GSM1014154) (Consortium, 2012; Davis et al., 2018; Thurman et al., 2012). These data are presented as differential profiles on one plot in Fig. 5B. Shaded area represents standard error in either direction for each 20-bp bin.

